# Selection of experience for memory by hippocampal sharp wave ripples

**DOI:** 10.1101/2023.11.07.565935

**Authors:** Wannan Yang, Chen Sun, Roman Huszár, Thomas Hainmueller, György Buzsáki

**Affiliations:** Center for Neural Science, New York University, NY, USA; Neuroscience Institute, NYU Grossman School of Medicine, New York University, New York, NY, USA; Mila - Quebec AI Institute, Montréal, Canada; Department of Psychiatry, New York University Langone Medical Center, New York, NY, USA

## Abstract

A general wisdom is that experiences need to be tagged during learning for further consolidation. However, brain mechanisms that select experiences for lasting memory are not known. Combining large-scale neural recordings with a novel application of dimensionality reduction techniques, we observed that successive traversals in the maze were tracked by continuously drifting populations of neurons, providing neuronal signatures of both places visited and events encountered (trial number). When the brain state changed during reward consumption, sharp wave ripples (SPW-Rs) occurred on some trials and their unique spike content most often decoded the trial in which they occurred. In turn, during post-experience sleep, SPW-Rs continued to replay those trials that were reactivated most frequently during awake SPW-Rs. These findings suggest that replay content of awake SPW-Rs provides a tagging mechanism to select aspects of experience that are preserved and consolidated for future use.

---

We recorded from many hundreds of neurons using dual-side silicon probes (Fig. 1A) implanted into the hippocampal CA1 pyramidal layer of mice (*1)*. To extract the sequential structure embedded in the spike data, we used sequence nonnegative matrix factorization (seqNMF) (*2*). seqNMF identified robust patterns that matched the behavioral events in the figure-8 maze (Fig. 1B-D; Suppl Fig. 1). Next, we used Uniform Manifold Approximation and Projection (UMAP) to embed the same high-dimensional data in a low dimensional space (*3*). UMAP visualization (Fig. 1E) revealed that population activity of the hippocampus corresponded to a latent space that topologically resembled the physical environment (*4, 5*). In addition, after color-labeling the manifold with trial numbers, we observed a prominent progression of states that corresponded to the trial sequence, spreading along one axis (Fig. 1F) (*6*). This observation was not dependent on the maze type or rodent species (Suppl Fig. 2) (*7*).

**Figure 1.**
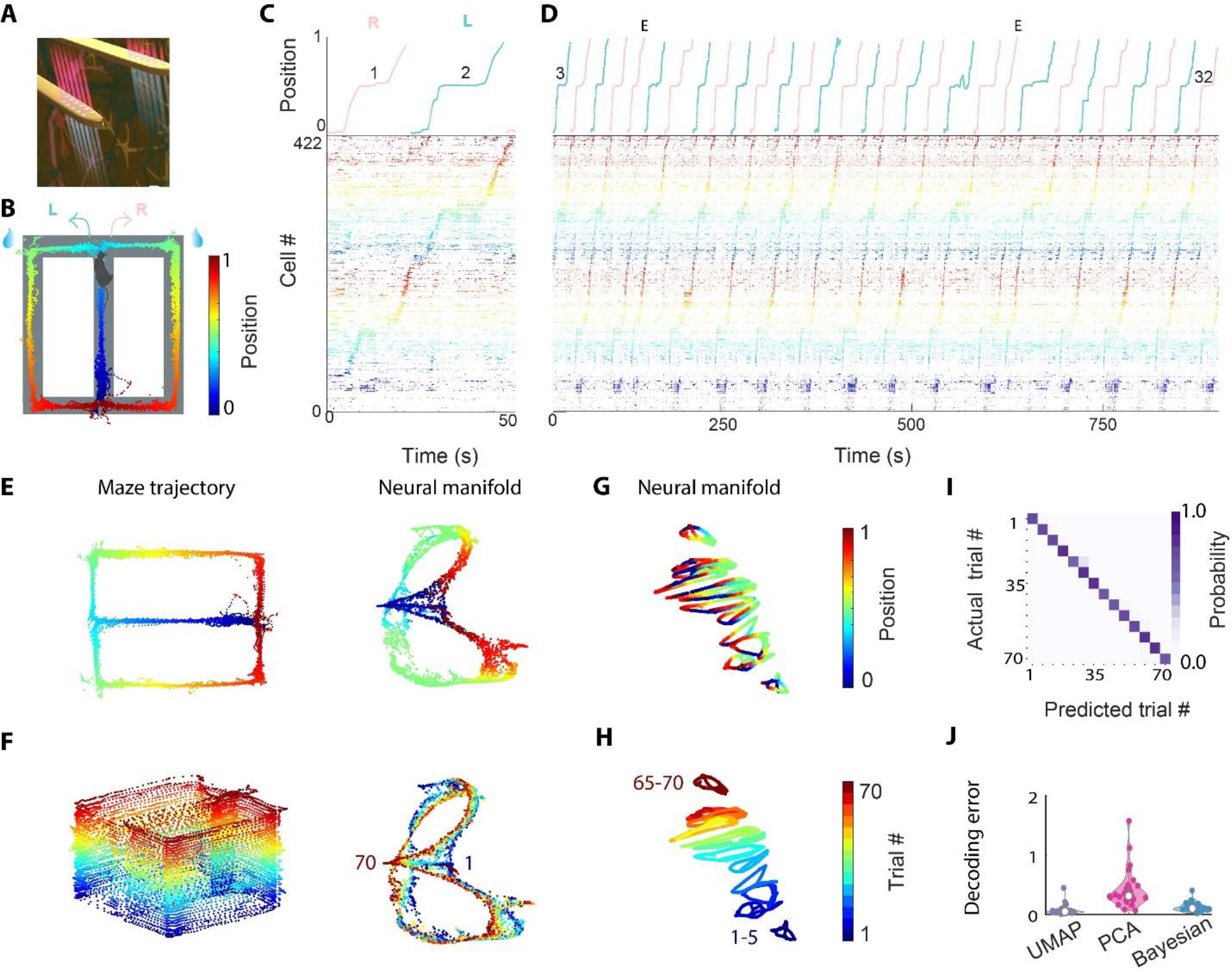
Trial identity can be decoded from the unique temporal evolution of neuronal population activity embedded in a low-dimensional manifold. **A**. Illustration of two shanks of the dual-sided probe. **B**. Figure-8 maze task, where mice alternated between left and right arms for water reward (blue droplets). The animal’s trajectory along maze corridors is color-coded by linearized position. **C**. Trials 1 (right) and 2 (left) during the figure-8 maze task. Top, linearized position of the animal. Red, right traversal; green, left traversal. Bottom, raster plot of spike sequences of 422 pyramidal cells simultaneously recorded from the right dorsal CA1 region, sorted by seqNMF and color-coded according to the linearized position, same color scheme as in Fig. 1B. **D**. Trial 3 to trial 32 (of 70 trials in total; Supp. Fig. 1) of an example session. E, error trials. **E**. Left, running trajectory of a mouse on the figure-8 maze. Right, UMAP embedding of population activity, colored by the mouse’s position. Each point corresponds to the low-dimensional representation of a binned spiking data from the high-dimensional space. **F**. Same session as E but the neural manifold is colored by trial number. **G**. Neural manifold of the same session as E, F, using semi-supervised UMAP trained on blocks of 5 trials. The manifold is colored by linearized position. **H**. Same as **G** but the neural manifold is colored by trial number. **I**. Confusion matrix of trial decoding accuracy based on UMAP embedding and KNN decoder (same session as in A-H) with ten-fold cross-validation. **J**. Trial identity decoding errors across all sessions using 3 different methods (Each dot in the violin plot indicates one session, pooled across n=26 sessions from 6 animals). Decoding error was measured in units of trial block where 5 trials were binned to 1 trial block. See also Suppl. Fig. 1-3.

To quantify the trial sequence information (*8*) present in the state space, we used the metric learning technique (see Methods) to generate the UMAP embedding in a semi-supervised manner. Although UMAP is typically used as a self-supervised dimensionality reduction tool. UMAP also can be used to learn a semi-supervised embedding (*3*), where labels are leveraged so that similar points are mapped closer together. Successive five trials composed a trial block, and a unique label was assigned to each trial block (Fig.1 G, H). Next, we tested whether trial-block membership of the unlabeled testing data could be decoded using a K Nearest Neighbor (KNN) decoder, followed by ten-fold cross-validation (Supp Fig. 5A). Trial identity could be accurately decoded from UMAP embedding (Fig. 1I). Those results were further confirmed using Principal Component Analysis (PCA) or Bayesian decoding (0.10, 0.42, 0.13, mean error for UMAP, PCA and Bayesian, respectively; Fig. 1J, Suppl Fig. 3), although these classical methods didn’t generate intuitive visualization for the initial hypothesis generation process (*9*).

To probe the contribution of the individual neurons to the population features, we first plotted the tuning curves of individual neurons across trials. We found diverse patterns of inter-trial variability: within-place field rate remapping (Suppl Fig. 4A, B) (*10*), place fields emerging on later trials (Fig. 2A bottom) (*11*) and shifting place fields across trials (Fig. 2A top; Suppl Fig. 4C) (*12*–*14*). In principle, decodability of trial identity could be explained by either stochastic or structured variation across trials. To identify the source of variability, we built a model that generated stochastic fluctuation in firing rate across trials. Each simulated cell’s spiking activity was generated through a Poisson process based on the tuning curve of real neurons (Fig. 2B; see Methods). We passed the simulated spike trains through the same dimensionality reduction pipeline as for the real data. As expected, the manifold of the simulated data also reflected the topology of the maze (Fig. 2C). In contrast, trial identity information across trials vanished (Fig. 2D-E). As an alternative to the standard ten-fold cross-validation method (Fig. 1K; Supp Fig. 5B, D), we implemented a targeted validation method (“leave one-trial out” procedure) that is expected to yield high decoding accuracy only if the state space of the data changed in a structured way, evolving along one axis according to the sequence of trial events. Since the training set did not contain any data that shared the trial identity of the testing set, the test set would decode only the closest trial in the state space (Suppl Fig. 5A). Indeed, we observed that only the real data were decoded correctly to the preceding or succeeding trials, occupying the space immediately next to the diagonal of the confusion matrix (1.20 and 7.84 mean error for real and simulated data for one example session; Fig. 2F; Suppl Fig. 5C). The trial decoding accuracy of real data was significantly higher than that of simulated data or trial-shuffle data (P < 10^-10^; 1.51, 4.07 and 5.34 mean error for real, simulated and shuffle data; Fig. 2G; Suppl Fig. 6). These findings indicate that inter-trial variation of population activity (drift) is highly structured. We also found that both place cells (PC) and non-place cells (NPC) contributed to trial identity decoding (0.1, 0.42, 0.49, 0.53, 0.59 for all cells, PCs, NPCs, PCs with size-matched control and NPCs respectively; Fig. 2H-I for ten-fold cross-validation results; Suppl Fig. 5E for hold entire-trial out results). The neural manifolds of different trials were aligned better with place cells than with non-place cells, possibly because place fields are consistent across trials. The trial identity decoding error was close to 0 for all sessions in the dataset (all sessions have >150 neurons), however decoding accuracy deteriorated rapidly when down-sampled to <100 neurons in a session, indicating that trial identity is encoded in the population (Fig. 2J; Suppl Fig.5 F for ten-fold cross-validation results; Suppl Fig. 7,8 for hold entire-trial out results).

**Figure 2.**
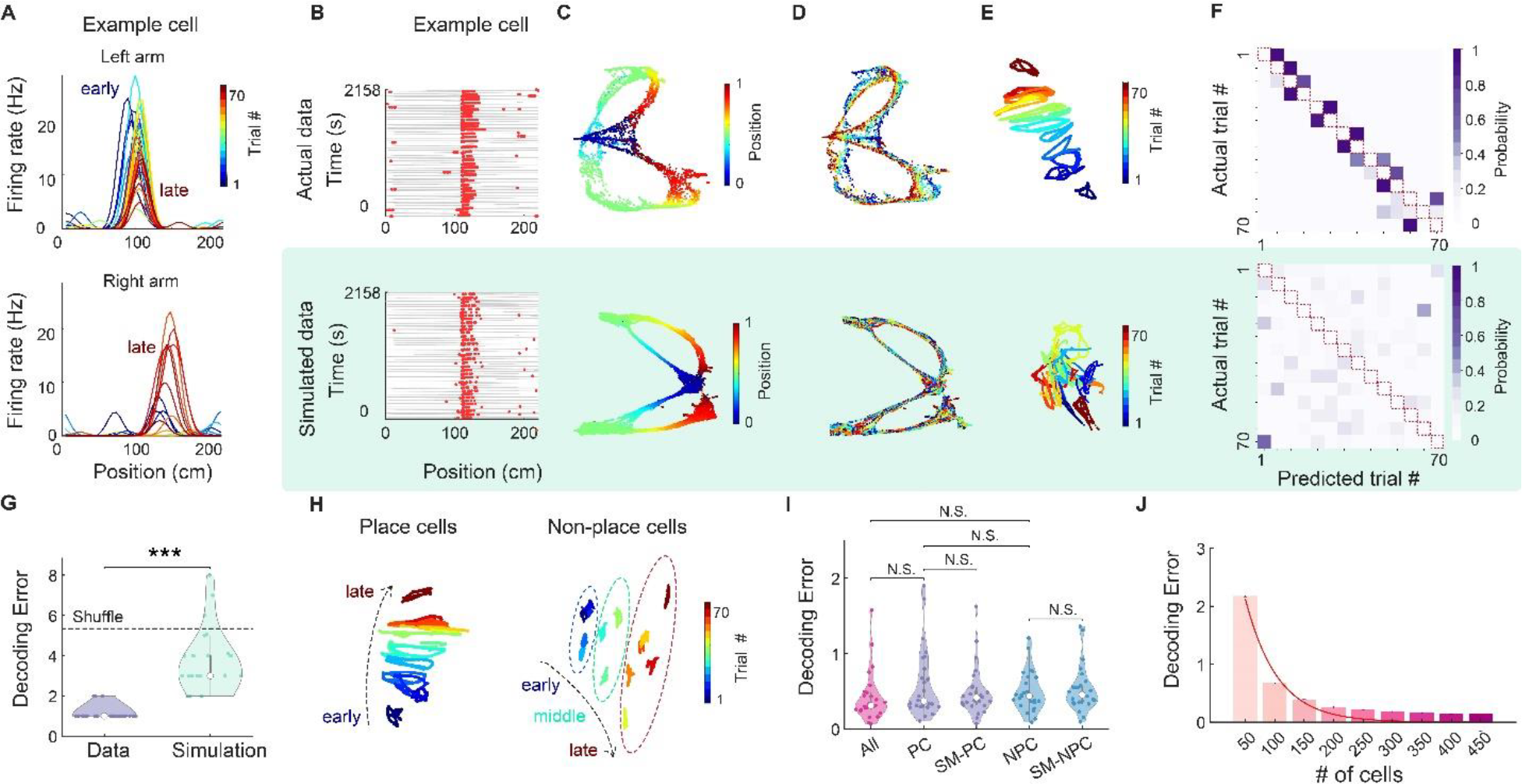
Contribution of single cell and subpopulation of cells to the structured trial identity coding. **A**. Trial-by-trial firing rate of an example neuron during left-arm (top) and right-arm trials (bottom) in the figure-8 maze. Color indicates the trial number. **B**. Raster plot of an example neuron (top) and a simulated neuron (bottom), generated from Poisson process based on the across-trial mean firing rate of the real neuron (see Methods). **C**. Top, UMAP manifold generated from the population activity of all the neurons in one example session, colored by linearized position. Bottom, UMAP embedding from the population activity of the corresponding simulation. **D**. Same as C but colored with the trial number. **E**. Neural manifold of the same session as C-D, using semi-supervised UMAP trained on blocks of 5 trials. **F**. Confusion matrix of trial decoding accuracy for real (top) and simulated population (bottom) using hold one-trial out cross-validation. Note that the diagonal (red dashed line) of the confusion matrix has 0 decoding probability, because the training and test data does not share data from the same trial block. **G**. Trial decoding error of real (purple) and simulated (green) data across all sessions (*** P < 10^-10^, un-paired t-test; n = 26 sessions from 6 animals). Dashed line indicates the chance level. **H**. Neural manifold generated from place cells (left) and none-place cells (right) from the same session as A-F, using semi-supervised UMAP trained on blocks of 5 trials. **I**. Trial decoding error of all cells (red), place cells (PC), and its size-matched control from all cells (SM-PC; purple), as well as non-place cells (NPC) and their size-matched control (SM_NPC; blue). Decoding error was measured in units of trial block where successive 5 trials were binned to 1 trial block. N.S., not significant, un-paired t-test; n= 26 sessions from 6 animals. **J**. Decoding error when downsampling the cells in the example session to subsamples of varying cell numbers (from 50-450 cells). Error bar indicates the standard error of mean (SE) across 1000 subsamples add statistics). Decoding error was measured in unit of trial block (where successive 5 trials were binned to 1 trial block). Red line, fitted exponential curve. See also Suppl. Figs. 4-8.

Place and trial sequence encoding is associated with theta frequency (6-9 Hz) oscillations (“theta state”) (*15*) as animals actively navigate in an environment. When animal stops to consume water reward, theta activity gives way to synchronous population events, known as sharp wave ripples (SPW-R; Fig. 3A, B). Since spike sequences within SPW-Rs are known to replay place field sequences (*16, 17*), we asked whether trial identity could also be detected in replay events. A candidate SPW-R event was deemed as significant replay when both its distance to manifold and trajectory length were significantly shorter compared to shuffle distributions (Fig. 3C, D; Suppl Fig. 9, 10, 11; See methods). The majority of SPW-Rs occurred in the reward area (Suppl Fig.12 A-C) and about 33% of the awake replays were significant replays (Fig. 3E) (*18*). We decoded not only the spatial trajectory but also trial identity of the significant replay events using different methods (PCA and UMAP) and quantified the trial-replay coherence within event as well as tested if decoding results match between different methods (Fig. 3F, G; Suppl Fig. 13 A-J). We found that the coherence of trial replay events (coherence between different decoding methods as well as among different time bins within the same event) was high but decreased rapidly as we down-sampled the number of cells used for decoding (Suppl Fig. 13 K, L). Next, we examined whether the SPW-Rs replayed the past, future, or current trial. Our results revealed that the spike content of SPW-Rs decoded most reliably the present trial (Fig. 3I) (*19*–*21*).

**Figure 3.**
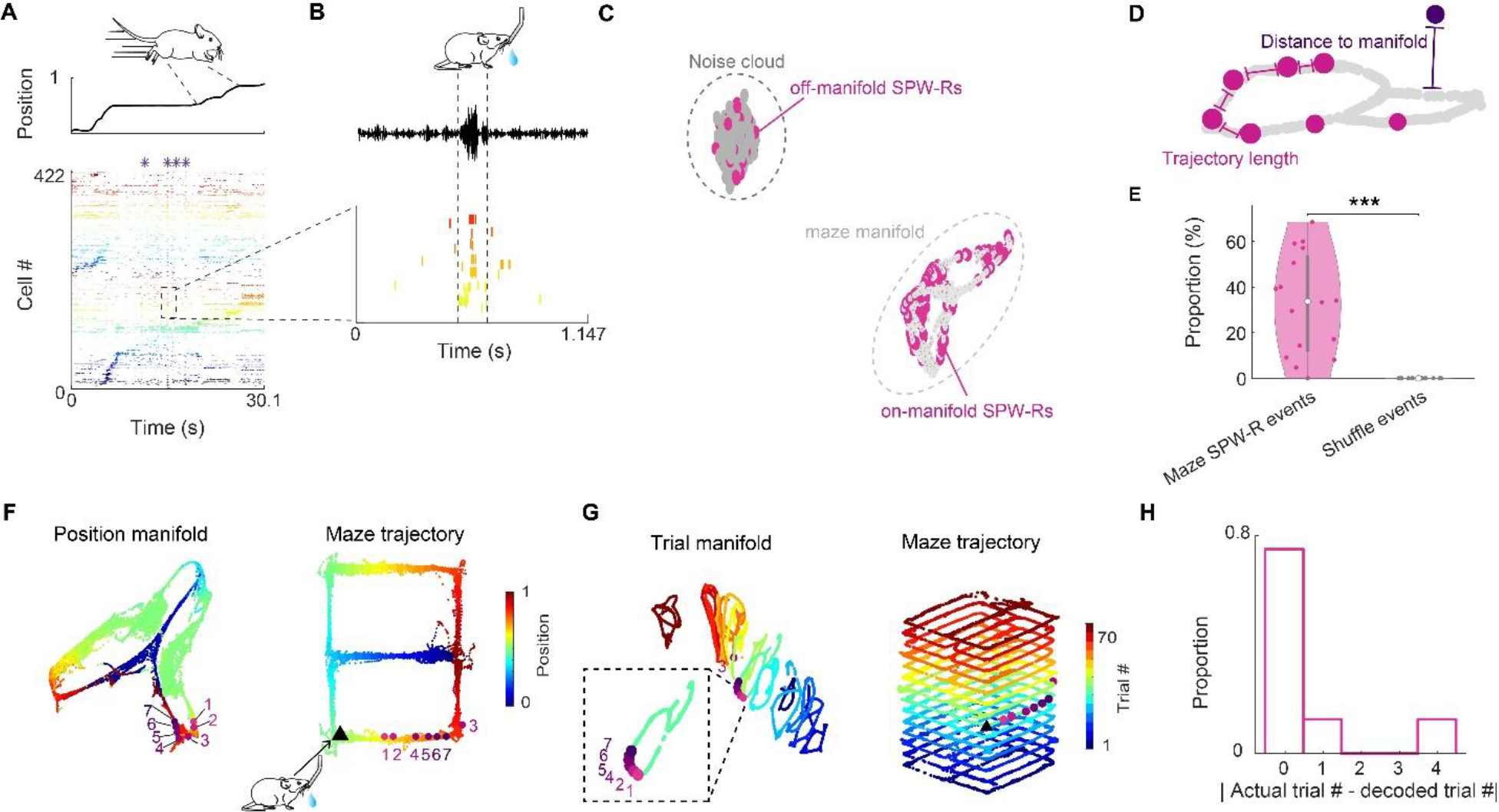
Maze SPW-Rs replay of trial events. **A**. Spiking activity of an example trial in the figure-8 maze. Top, linearized position of the mouse. Bottom, raster plot of neuronal spiking activity sorted by seqNMF. Purple stars, SPW-Rs. **B**. Zoom-in display of a replay event. Bottom, raster plot of neurons belonging to a sequence factor with significant reactivation strength (see Methods). Neurons were sorted in the same order as in A. Top, LFP filtered in the ripple band. **C**. All awake SPW-R events (red) in this example session were embedded with the neural manifold during navigation (light gray) and the noise cloud consisted of negative samples (dark grey). Two clusters of SPW-R replay events were distinguished: Off-manifold and on-manifold events. (See also Supp. Fig. 9, 10). **D**. SPW-R replays were classified as significant when they occurred near the maze manifold and when their trajectory length along the manifold was shorter in comparison to shuffle (see Methods; Suppl Fig. 9, 11). **E**. Percentage of significant replays in the maze (# of maze significant replay events / total # of maze SPW-R events) compared to shuffle. *** P < 10^-4^; un-paired t-test; n = 26 sessions from 6 animals. **F**. UMAP embedding and decoded position for the same event as in panel B. Left, the neural manifold during maze running (‘position manifold’, colored by position). The pink-purple dots represent the neural embedding of 7 successive time bins of a SPW-R replay event (each time bin is 20 ms); Right, the replay content of each time bin was decoded to a position bin along the maze trajectory according to the position label of its nearest neighbor on the manifold. Black triangle, physical location of the mouse when the replay event took place. **G**. Left, the same event was embedded with the ‘trial manifold’. Right, replay content of each time bin was decoded as the trial block label of its nearest neighbor on the manifold (each trial block corresponds to 5 trials). **H**. Distribution of differences between the actual trial block of SPW-Rs events and their decoded trial block identity across all sessions. See also Suppl. Figs. 9-13.

To examine the relationship between awake SPW-Rs and those during post-experience sleep, we compared the population activity of awake SPW-Rs in the maze with that during post-experience sleep (post-sleep) in the home cage (Fig. 4A-E). We observed that the distribution of decoded trial identity during post-experience sleep was highly correlated with that during maze replay, but not with pre-experience sleep replay (R = 0.86, P <10^-36^ for correlation between post-sleep replay and maze replay; Fig. 4F-H; Suppl Fig. 14A, B) (*23, 24*). We compared which factor best explained the trial distribution pattern during post-sleep, using a mixed-effect linear regression analysis. The strongest predictor for the trial distribution pattern during post-experience sleep was that of the SPW-R replay in the maze (P < 10^-26^, P = 0.06, P < 10^-4^, P < 0.89, P = 0.87 for in-maze replay, theta power, theta cycle count, pre-sleep replay, and shuffle, respectively; Fig. 4I) (*22*). The same conclusion was held when we conducted the same regression analysis after using PCA for dimensionality reduction (Suppl Fig. 14C).

**Figure 4.**
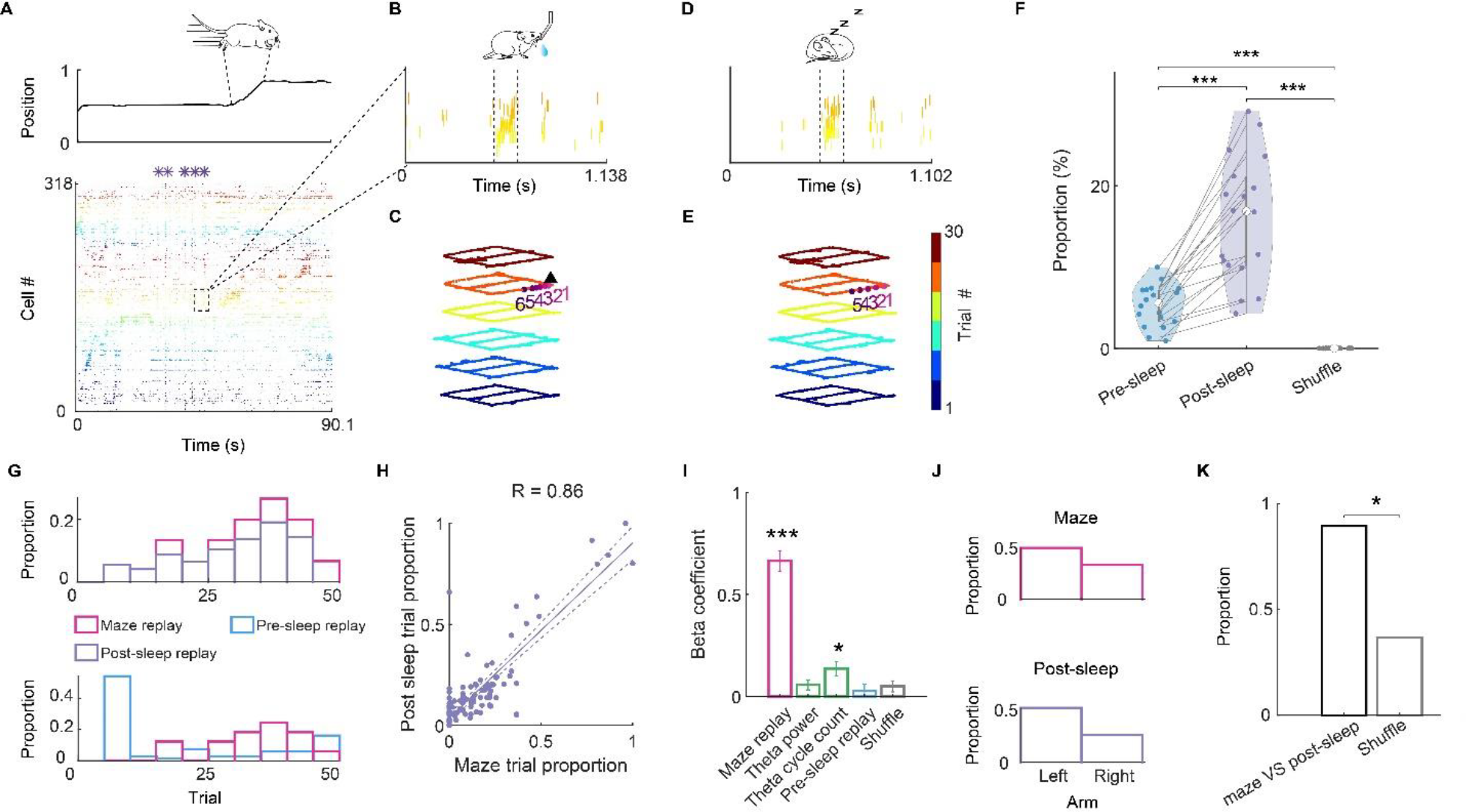
Replay of trial identity and maze segments during sleep can be predicted from awake SPW-Rs. **A**. Spiking activity in the figure-8 maze. Top, linearized position of the mouse. Bottom, raster plot of neuronal spikes, sorted by seqNMF. Purple stars, SPW-Rs. **B**. Zoom-in display of a wake replay event in the maze. The raster plot contains neurons belonging to the sequence factor with significant reactivation strength (see Methods). Neurons were sorted in the same order as in A. **C**. decoded position and trial identity of successive 20-ms bins (1 to 6 bins) of the same SPW-R replay event in B. Black triangle, physical location of the mouse when the replay event took place. **D**. Raster plot of a SPW-R replay event during sleep in the home cage. **E**. decoded position and trial identity of successive 20-ms bins (1 to 5 bins) of the same SPW-R replay event in D. **F**. Percentage of significant replays during pre-experience and post-experience sleep in the home cage, compared with shuffled data. Pre-sleep vs post-sleep, *** P < 10^-5^; post-sleep vs shuffle, *** P < 10^-8^; pre-sleep vs shuffle, *** P < 10^-8^; un-paired t-test. n = 26 sessions from 6 animals. **G**. Distribution of the replay events across trialss decoded from the population spike content of SPW-Rs in an example session. Top, maze and pos-sleep replay trial distribution pattern. Bottom, maze and pre-sleep replay trial distribution pattern. **H**. Correlation between trial distributions of trial blocks during maze and post-sleep SPW-Rs replay. Pearson Correlation, R = 0.86, P <10^-34^; n = 16 sessions from 5 animals. **I**. The predictive relationship between trial distribution pattern of post-sleep SPW-R and other candidate variables, including the trial distribution patterns of theta cycle, theta power, pre-sleep, and trial-shuffle data (refer to Suppl Fig. 6 for procedure for generating trial shuffle data). *** P < 10^-23^ for awake replay; *** P < 10^-3^ for theta cycle number; The relative predictive power of a given metric was considered non-significant when it overlapped with zero. n = 16 sessions from 5 animals. **J**. Proportion of maze segment replays (left arm and right arm) during awake and post-maze sleep SPW-Rs in an example session. **K**. Proportion of sessions with same rank order between maze segments during awake and post-maze sleep SPW-R replays. *P < 0.05, Chi-square test; n = 13 sessions from 5 animals. See also Suppl. Figs. 14.

Finally, we examined the relationship between awake and sleep replays from a different angle, we next examined how replays of the left versus right arms during wake and post-sleep SPW-Rs were correlated, exploiting the natural variability in the distribution of maze arm replays in different sessions (Fig. 4J; Suppl Fig. 12A, B, D). The correlation of maze arm replays between wake and sleep SPW-Rs was significantly higher than in the shuffled data (Fig. 4K). The difference in replay proportion across left and right arms could not be explained by the difference in the decidability (measured by spatial information) nor the difference in coverage (measured by number of visits) between the two arms (Suppl Fig. 14D-G).

Exploration of the environment is a regular alternation between ambulation and rest phases, enabling brain state changes (*23*–*25*), and replaying aspects of experience during SPW-Rs (*16, 17, 20*). We hypothesize that awake SPW-Rs represent a natural tagging mechanism of experiences (*26*). In turn, the uniquely tagged neuronal patterns are reactivated numerous times during SPW-Rs of post-task sleep to consolidate the selected experience and combine it with the existing knowledge base of the brain (*27, 28*).

Previous observations support this scenario. Cues relevant to future behavioral success can bias the neuronal trajectories of SPW-Rs (*29*–*31*). Perturbation of SPW-Rs prevents place field stabilization (*32*) and experience to be (*33*–*39*). In humans, items whose electrophysiological brain patterns are not reactivated during non-REM sleep are forgotten (*40*). Replay can occur after a single experience and the number of SPW-R after learning predicts subsequent memory performance (*21, 38, 41*–*43*).

Salient features of the experience, such as novelty (*44*–*47*) and reward (*48*), increase the incidence of SPW-Rs and replays (*49, 50*). In our experiments, we could exclude differences in those external factors as alternative reasons for shaping sleep replay patterns by demonstrating uniquely changing neuronal population activity patterns within the same maze and session. Selective tagging of trial events by awake SPW-Rs could occur because the perpetually evolving population activity pattern during theta state during theta state can serve as a temporal scaffold to organize experiences (*51*), and differentiate successive events.

Our observations suggest that reward provides an affordance for shifting brain states and it is the awake SPW-R that serves as a mechanism for selection (*41, 52*). These findings establish a neurophysiological framework for multiple domains of systems neuroscience, including credit assignment (*16, 28*), representational drift (*8, 51*), “event remapping” (*8*), time coding (*6, 53*–*55*), and memory editing (*56*).

## ACKNOWLEDGEMENTS

We thank Kirill Kiselev, Kanishk Tewatia and Julia Paraiso for helping solve technical problems with data visualization and validate ripple detection results. We thank Andreas Grosmark for sharing his data. We thank Ipshita Zutshi, Kathryn Mcclain, Zheyang (Sam) Zheng and all members of the Buzsaki lab for their insightful discussions and feedback. We greatly appreciate David Tingley, Jo Carpenter and Kiah Hardcastle for their constructive feedback on the manuscript. This work was supported by NIH grants (R01MH122391; U19NS107616).

## AUTHOR CONTRIBUTIONS

Conceptualization: WY, GB

Methodology: WY, RH, CS

Investigation: WY, RH, TH

Visualization: WY, CS

Funding acquisition: GB

Project administration: GB

Supervision: GB

Writing – original draft: WY, CS, GB

