## SUPPLEMENTARY MATERIAL for "Selection of experience for memory by hippocampal sharp wave ripples"

### METHODS

#### Implantation and recording.

The data in this paper came from 3 different datasets. Dataset 1 was the main dataset used for both visualization and replay decoding. Only 1 session was taken from dataset 2 and 3 each, and the results were presented only in supplementary figure 2 for visualization. The purpose of the visualization is to demonstrate that the trial specific population activity pattern can be observed in diverse tasks and different rodent species.

Dataset 1: As described in (*1*), chronic recordings were performed from freely moving adult C57BL/6J mice. The mice were anesthetized with 1.5–2% isoflurane and implanted with high-density silicon probes, mounted on a microdrive. The ASSY Int64-P32-1D or ASSY Int128-P64-1D silicon probes (Diagnostic Biochips) were dual-sided designs, equipped with recording sites on both sides of each shank. The spacing between shanks was 250 µm. Craniotomy was drilled at −2mm anteroposterior (AP) and 1.7mm mediolateral and the probe was lowered to the deep neocortical layers, and the drive was cemented to the skull. A stainless-steel screw was placed over the cerebellum for grounding and reference. Craniotomies were sealed with a mix of dental wax and mineral oil, and a copper mesh cage was constructed to provide electrical and mechanical shielding. Postoperatively, animals received a single intramuscular injection of 0.06mg/kg of buprenorphine (0.015mg/ml) and as needed for the next 1–3 d. Animals were allowed a 7-d recovery period before the start of experiments. After a 7-d recovery period, neural signals were recorded in the homecage while probes were lowered into the CA1 pyramidal layer, which was identified physiologically via the sharp wave polarity reversal. Neural data were amplified and digitized at 30 kHz using Intan amplifier boards (catalog no. RHD2132/RHD2000, Evaluation System).

Dataset 2: As described in (*7*), male Long-Evans rats (250-350g) were bilaterally implanted in the dorsal hippocampus with two 8 shank (BUZ64) silicon probes. Each shank of the 8-shank silicon probes had 8 sites. All sites were vertically staggered along the shank with 20 µm spacing between sites. Each site had an area of 160 µm2 and an impedance of 1–3 MΩ. In each rat, 50 µm wires were placed gently abutting the left, right mastoid masseter as well as the back neck muscle for electromyographic (EMG) recordings used in sleep classification. All silicon probes were implanted parallel to the septo-temporal axis of the dorsal hippocampus. Finally, each rat was fitted with a small 3-dimensional accelerometer to record the animals' movement, or its absence, during sleep. Two stainless steel screws implanted above the cerebellum were used for referencing and grounding. Implantation was performed under isoflurane (1-1.5%) anesthesia. Each silicon probe was attached to a micromanipulator and lowered over the course of several days until hippocampal layer CA1 was reached as determined by the appearance of hippocampal CA1 sharp wave ripples and pyramidal cell activity. Probe placement was histologically confirmed *post hoc*.

Dataset 3: Procedures were identical to those for dataset 1 with the following exceptions: A 8-shank, single sided 128 channel probe (128-8, Diagnostic Biochips) was implanted transversely into the right dorsal hippocampus (medial edge 2.2 mm posterior, 1.1 mm lateral of bregma, lateral edge 1.5 mm posterior and 1.8 mm lateral of bregma). Peri- and postoperative analgesia was performed with subcutaneous Ketoprofen 10 mg/kg daily for 3 days.

**Behavior.**
Dataset 1: Mice were trained on a spatial alternation task in figure-eight maze and linear maze (*1*). Animals were water restricted before the start of experiments and familiarized with the maze (Fig. 1b) raised 61 cm above the ground. Over several days after the start of water deprivation, animals were shaped to visit alternate arms between trials to receive a water reward in the first corner reached after making a correct left/right turn. A 5-s delay in the start area was introduced between trials. Infrared (IR) sensors were used to detect the animal's progression through the task and 3D printed doors mounted to servo motors were opened/closed to prevent the mice from backtracking. IR sensors and servo motors were controlled by a customized Arduino-based circuit. The position of head-mounted red LEDs (light-emitting diodes) was tracked with an overhead Basler camera (catalog no. acA1300-60 gmNIR, Graftek Imaging) at a frame rate of 30Hz, and tracking data were aligned to the recording via transistor–transistor logic pulses from the camera, as well as a slow pulsing LED located outside the maze. In all sessions that included behavior, animals spent ~120min in the homecage before running on the maze and another ~120min in the homecage afterward. All behavioral sessions were performed in the mornings (start of the dark cycle). Only sessions with more than 150 cells and >20 trials were selected for analysis. n = 26 sessions from 6 animals (21 out of 26 sessions were figure-8 maze sessions, and 5 of them were linear maze sessions).

Dataset 2: As described in (*7*), a rat (Achilles) was extensively handled both before and after surgery. Water restriction was initiated one week after the surgery; animals were restricted to 90% of their initial body weight and given one day a week of *ad lib* water access. The rat was well acclimatized and recorded in the 'familiar' room (where all sleep recordings were performed) for at least one week prior to novelty maze sessions. This time was used to gradually lower the silicon probes into position. All hippocampal, EMG and accelerometer signals were recorded continuously at 20 kHz using four identical 256-channel Amplipex Systems (16-bit resolution; analog multiplexing; one in the familiar room and one in each of the two novelty rooms). Only one session (Achilles_10252013) from this dataset was used to produce the visualization in Suppl. Fig. 2.

Dataset 3: The mouse was handled and familiarized with the maze for multiple days prior to commencement of experiments. It was then trained to forage for rewards on the radial 8-arm maze (*54*). The maze consisted of eight concentrically arranged arms of hard plastic (each arm 40 cm long) with 5 cm high side walls that were elevated 30 cm above the floor. In our version of the task, three out of the eight arms of the maze were chosen semi-randomly every day, with the constraints that a maximum of two adjacent arms were rewarded and that at least 2 arms had to be different from those of the previous day. These three arms were initially 'baited' with water rewards (they were released through a solenoid valve that was automatically triggered when the animal crossed an infrared barrier in the middle of the arm) but only re-plenished after the animal had visited all rewarded arms. There were no discrete trials and the animals were allowed to continuously forage the maze for 20 minutes. The animal was then placed in its home cage for 2 hours, during which neuronal activity was recorded. After this, the animal was given another 20-minute session in the radial arm maze to assess for memory retention. Reward locations were kept constant between the two sessions of a given day but changed between subsequent days. Electrophysiological and behavior recording procedures were identical to those outlined for dataset 1 above. Only one session (TH11_210605) from this dataset was used to visualize position coding by UMAP of a radial arm maze (Suppl. Fig. 2).

**Unit isolation and classification.** Spikes were extracted and classified into putative single units using KiloSort1 and manual curation was performed in the Phy2 software with the aid of customized plugins (https://github.com/petersenpeter/phy2-plugins), as described in ref. (*1*).

**Place field and spatial information.** Place field analysis was done as ref. (*55*). The code is available at https://github.com/valegarman/HippoCookBook.The firing rate distribution within place fields ('rate map') was generated by first binning spiking data into 5 cm wide bins, and spike counts were normalized by time occupancy (smoothing size: 2 bins). Trials for forward and backward directions were evaluated separately. Place fields were defined from these rate maps as previously described (*35*). The field boundaries were defined when the rate decreased below 20% of the peak firing rate. Place fields cleared all criteria when they were between 8.75 cm and 75 cm wide, had a minimum peak firing rate of 3 Hz and a spatial coherence > 0.7 (*35*)

Spatial information was calculated in bits per spike as following:

$$SPI = \sum_{i}^{N} pi \frac{\lambda_i}{\lambda} log_2 \frac{\lambda_i}{\lambda} \quad (1)$$

where $\lambda_i$ is the mean firing rate of a unit in the $i^{th}$ bin, $\lambda$ is the overall mean firing rate and $pi$ is the probability of the animal being in the $i^{th}$ bin (occupancy in the $i^{th}$ bin/ total recording time).

**Poisson process for modeling single cell activity across trials.**
The simulated spike was modeled based on a Poisson process:

$$Pr(X = \kappa) = \frac{\lambda^{\kappa} e^{-\lambda}}{\kappa!} \quad (2)$$

The procedure to generate Poisson spikes was followed by (*56*). For each simulated neuron, the spike train progresses through time in small windows of ***dt (1ms)***, and spikes were generated via Poisson process using the cross-trial mean firing rate $fr(pos)$ of the real neuron at position ***pos***.

**Theta-cycle detection**. Theta cycle detection was done following ref (*55*). As theta-phase shifts along the radial axis, a channel with a positive sharp wave (that is, above the center of the pyramidal layer) was selected to ensure consistency of extracted phases across recordings. Broadband LFP was bandpass filtered between 6 and 12Hz using a fourth-order Chebyshev filter. The Hilbert transform of the filtered signal was computed, and its absolute value and angle at each timepoint were taken to be the theta-band amplitude and phase, respectively. Intervals with theta-band amplitude 1 s.d. above the mean were considered for theta cycle detection. Within these intervals, timepoints where the phase crossed 0° were identified as peaks of theta, and timepoints of consecutive theta peaks were considered the onsets and offsets of individual theta cycles (all throughout, peaks are at 0° and 360° and troughs at 180°). Only theta cycles occurring within identified waking periods (see section State scoring) were considered for analysis.

**State scoring**. State scoring was performed as described previously (https://github.com/buzsakilab/buzcode/blob/dev/detectors/detectStates/SleepScoreMaster/SleepScoreMaster.m). First, the LFP was extracted from wideband data by low-pass filtering (sinc filter with a 450-Hz cut-off band) and downsampling to 1,250Hz. Three signals were used for state scoring: broadband LFP, narrowband theta frequency LFP and electromyogram (EMG). Spectrograms were computed from broadband LFP with fast-Fourier transform in 10-s sliding windows (at 1 s) and principal component analysis (PCA) was computed after a z transform. The first PC reflected power in the low (<20-Hz) frequency range, with oppositely weighted power at higher (>32-Hz) frequencies. Theta dominance was quantified as the ratio of powers in the 5 to 10-Hz and 2 to 16-Hz frequency bands. EMG was estimated as the zero-lag correlation between 300 and 600 Hz filtered signals across recording sites. Soft sticky thresholds on these metrics were used to identify states. Briefly, high LFP PC1 and low EMG were taken to be NREM, high theta and low EMG were considered to be REM and the remaining data were taken to reflect the waking state. All assignments were inspected visually and manually curated wherever appropriate(https://github.com/buzsakilab/buzcode/blob/dev/GUITools/TheStateEditor/TheStateEditor.m).

**Ripple detection.** Ripple events were conditioned on the coincidence of both population synchrony events, and LFP detected ripples as described in ref. (*7*). For each session the combined spiking of all recorded CA1 pyramidal cells were binned in 1ms bins and convolved with a 15 ms Gaussian kernel. For each session a trigger rate was defined as being 3 standard deviations above the mean of all 1 ms bins within NREM epochs of both PRE and POST epochs combined. Putative population synchrony events were detected when the smoothed firing rate vector crossed the trigger rate. The beginning and end of the putative synchrony events were defined as the time points at which the convolved firing rate vector returned to the mean of all within-NREM firing rate bins. Independently, ripple events were detected from the pyramidal layer LFP. Population synchrony events that did not contain at least one LFP-detected ripple were discarded Population synchrony events which (1) contained at least one LFP-detected sharp wave-ripple, (2) lasted between 50 to 500 ms, and (3) occurred during non-theta or 'off-line' states (quite waking [immobility] or NREM) and 4) in which at least five distinct pyramidal cells each fired at least one spike were termed 'Ripple events' and considered for further analysis. This 'five pyramidal cell' inclusion criterion was used as it is the minimum number of cells needed to establish significance for sequence relationship between cells at $p < 0.05$ (5! = 120 possible orderings, of which half are mirror 'forward' and 'reverse' sequences and hence equivalent with respect to the sequence analysis, resulting in 60 unique orderings, 1/60 = 0.0167).

**SeqNMF.** SeqNMF algorithm was implemented following ref. (*57*) and code from (https://github.com/FeeLab/seqNMF). Briefly, seqNMF started by taking a data matrix X which contains the activity of N neurons at T time points. If the neurons exhibit a single repeated pattern of synchronous activity, the entire data matrix can be reconstructed using a column vector w representing the neural pattern and a row vector h representing the times and amplitudes at which the pattern occurs. Data matrix X was mathematically reconstructed as the outer product of w and h. If multiple component patterns are present in the data, then each pattern can be reconstructed by a separate outer product, where the reconstructions are summed to approximate the entire data matrix as follows:

$$X_{nt} \approx \sum_{k=1}^{K} W_{nk} H_{kt} \ = \ (WH)_{nt} \qquad (3)$$

Where $X_{nt}$ is the ***(nt)****th* element of matrix **X**, which represents the activity of neuron n at time t. In order to store K different patterns, **W** is a N x *K* matrix containing the *K* exemplar patterns or sequence factors (Suppl Fig. 1A) and **H** is a *K* X T matrix. Here we choose *K*= 8, given the topology of the figure 8 maze.

**Low dimensional manifold visualization with UMAP (unsupervised).** Neural data were first preprocessed before dimensionality reduction. Neural spiking data (spike count) during maze learning was binned into 100ms bin. The data was then smoothed using a 500 ms wide Gaussian kernel. The nonlinear dimensionality reduction algorithm UMAP was then applied to this matrix. Each point in the low dimensional manifold corresponds to the population activity at a single time bin in the session, and collectively the cloud of points maps out the animal's navigational trajectories during the task. The code is available at https://github.com/lmcinnes/umap/blob/master/doc/how_umap_works.rst.
UMAP hyperparameters: n_neighbors = 20, metric = 'cosine', output_metric = 'euclidean', learning_rate = 1.0, init = 'spectral', min_dist = 0.1, spread = 1.0,repulsion_strength = 1.0, negative_sample_rate = 5, target_metric = 'categorical', dens_lambda = 2.0, dens_frac = 0.3, dens_var_shift=0.1.

**Low dimensional manifold visualization with UMAP (supervised**). There are two UMAP procedures: 1. Unsupervised dimensionality reduction. 2. Supervised (and semi-supervised) dimensionality reduction, which makes use of categorical label information. With supervised methods (*3*), data from the same trial were given the same supervision label. Data points that share the same labels are leveraged so that similar points are embedded closer together than they would otherwise be. For example, visualizations in Fig. 1G, H were produced by supervised UAMP dimensionality reduction, where we exploited this option by using trial identity labels in the behavioral data as supervised information. This was implemented by passing the trial block number as target when calling the fit_transform function from the UMAP python package https://github.com/lmcinnes/umap/blob/master/doc/how_umap_works.rst.

**Decoding maze data with UMAP.** There were 4 steps for UMAP decoding: (1) Data preprocessing; (2) Manifold embedding with training data; (3) Embedding testing data based on the learned metric generated from training data and (4) K-nearest neighbor (KNN) decoding. Neural data were first preprocessed before dimensionality reduction as described above.

To decode the trial identity from population activity, metric learning was implemented (*3*). Briefly, metric learning is the process where a labeled set of points was used to learn a metric on data, and the learned metric was used as a measure of distance between new unlabeled points. In our case, a subset of data was selected as training data, based on which the manifold embedding was generated. The training data was supervised with trial labels by concatenating 5 trials as one block. After generating the manifold with training data, previously unseen (and unlabeled) testing points were then embedded into the learned space using the function in the Python package (https://umap-learn.readthedocs.io/en/latest/supervised.html).

The scikitlearn implementatio of K-Nearest Neighbors (KNN) algorithm (https://scikit-learn.org/stable/modules/generated/sklearn.neighbors.KNeighborsClassifier.html) was used to decode the trial identity of the testing data embedded in the low dimensional manifold. Uniform weights were applied on distances to 10-nearest neighbors. Since the lower dimensional UMAP embeddings are obtained with Euclidean metrics, we used Euclidean distance metrics for KNN as well. The mode of the 10-nearest neighbors of each time point was taken as the decoded trial of each time bin. Position identity was decoded following an analogous procedure, the only difference was that the training labels were position bin (5-cm bin) instead of trial block.

**Decoding error and validation method.** Decoding error during behavior was cross-validated by tenfold cross-validation, which was implemented by splitting the data into 10 subsamples, each randomly drawn from across the entire session. Decoders were then trained on 9 of the subsamples and tested on the remaining subsample. All 10 subsamples were used as test data during successive iterations. Decoding error was taken as the average error across all 10 trained decoders. For PCA decoding, cross-validation was repeated 1,000 times with overall decoding accuracy taken as the mean across the 1,000 repetitions. Because of computational constraints, cross-validation was repeated 10 times when using UMAP (Suppl Fig. 4A).

For the purpose of testing if the state space of different trials evolved systematically along a single axis whereby neighboring trials were embedded closer together in the neural manifold, a targeted validation method was also implemented. In contrast to tenfold cross-validation, an entire trial block was held out for each subset of training data. Since the training data does not contain any data that share the same trial identity as the testing data, testing data was expected to be decoded to the neighboring trial only if the population activity of neighboring trials is more similar than that of distant trials (Suppl Fig. 4B). Statistical significance for decoding error was determined by comparing mean decoding error from the original data against mean decoding error from trial shuffled data. Trial shuffled comparisons were generated by shuffling the trial identity of the data. This process was repeated 1,000 times.

For comparing decoding accuracies across different subpopulations of place cells and none-place cells, size-matched populations were used as control. Size-matched populations were generated by subsampling the total population of cells for a given session without replacement.

For downsample analysis, different subsets of cells were randomly drawn from the total population of cells for a given session without replacement. For each sample size, 1000 different random seeds were used to randomly subsample different cells. Decoding accuracy was taken as the average accuracy across all 1000 different random samples.

**Decoding SPW-R content with UMAP**. The flow diagram illustrating the procedures for SPW-R events decoding was shown in Suppl Fig.9. Step 1: Preprocessing data. Spike trains during maze learning were binned in 100 ms time bins and then smoothed. Spike trains of candidate SPW-R replays were binned in 20ms time bins. Step 2: generating manifold embedding using unsupervised manifold learning (UMAP). The preprocessed data matrix was then passed through a nonlinear, dimensionality reduction step by UMAP, which generated a three-dimensional nonlinear embedding, allowing the topological structure to be visualized. This step is unsupervised. For further details related to step 1 and 2, refer to Supl. Fig. 10. Step 3: Projecting candidate ripple events to the manifold generated at step 2. Step 4: measuring event distance to manifold and trajectory length. The event distance to manifold is defined as the mean Euclidian distance to manifold across all time bins within the event. Step 5: Testing if the candidate events were significant according to two criteria. (1) mean *distance* to manifold of a candidate event needs to be significantly smaller than shuffle (shuffle cell identity). (2) The second criterion is that the mean trajectory length needs to be significantly smaller (i.e. no large stochastic jumps on the manifold) compared to shuffle (shuffle time). Refer to Supl Fig. 11 for the corresponding details of how the shuffle distributions were generated. Step 6: If a candidate event is a significant replay event, proceed to decode both the position and trial identity of the replay event. To decode the position relayed by the candidate event, the population activity during ripple was projected to the position manifold where each point on the manifold is associated with a position bin label (positions were binned into 5-cm bins). The low dimensional embedding along with the position label were used to train a KNN decoder. Replay content of each time bin of a candidate event is decoded by taking the mean position bin of its K nearest neighbors on the position manifold. Similarly, the ripple activity vectors were projected to the trial manifold for trial identity decoding. The trial manifold was learned through semi-supervised dimensionality reduction (trials were binned into 5-trial bins).

**UMAP and negative sampling**. UMAP took a sampling-based approach called negative sampling (*3*), a contrastive method, to optimize the low dimensional embedding. Briefly, the intuition behind negative sampling is that data far away from the original high dimensional space is sampled as negative samples from which data sample is repulsed away from, whereas similar data in the high dimensional space are pulled closer to each other. The shuffle data serve as negative samples to generate the correct embedding for testing data, such that population vectors similar to the maze manifold will be pulled closer to the maze manifold in the low dimensional space and those dissimilar to the maze manifold will be pulled closer to the shuffle data cloud. Without the presence of shuffle data, all SPW-R data would have no choice but be incorrectly embedded to the maze manifold (Suppl Fig.10 A-D). With the shuffled noise cloud, the population vector of a certain time point was 'attracted' to the behavior manifold if it is close enough, or to the noise cloud if it is not (Suppl Fig.10 E-H). Ripple shuffle data almost exclusively attracted to the noise cloud (Suppl Fig.10 I-J).

**Elaboration on the criteria on significant replay**. Candidate event was binned into 20ms bins. we measured Euclidean distance of each data point (since the distance metric selected for embedding in low dimensional space with UMAP is Euclidean distance) to the k nearest neighbor on the maze manifold. Two kinds of shuffles were used to generate null distribution for distance to manifold and trajectory length respectively. (1) *Cell identity shuffle* (Suppl Fig. 11B). The purpose of cell identity shuffle was to degrade population pattern without affecting trajectory length and other properties such as firing rate and firing rate variance of each neuron. (2) *Temporal shuffle. It* specifically degrades trajectory length of the event while preserving the distance to manifold (Suppl Fig. 11A). A replay event was classified significant only when it was significantly different from both null distributions generated from the two shuffle methods (Suppl Fig.11 C, D).

**Decoding with PCA and KNN algorithm**, KNN was applied to the low dimensional data after linear dimensionality reduction with PCA. To decode position information, training data labeled with position was used to train the KNN classifier. The position of the testing data was then decoded based on their distance to the k nearest neighbor in the training set. Similarly, trial block identity was used as training labels for trial membership decoding.

**Decoding with a Bayesian decoder.** For Bayesian decoding, neural spiking data was preprocessed in a similar way for UMAP decoding. The data was then smoothed using a 1000ms wide Gaussian kernel. Only data with speed larger than 2-5cm/s was taken to build the template. Bayesian decoding was performed independently for the left and right arm trial. For each ripple event, the position and trial identity of the event was decoded separately.

**Bayesian position decoding.** Bayesian decoding of position was performed as described in ref. (*7*). Briefly, for each of the binned firing rate vectors, Bayesian classification of virtual position was performed utilizing a 'template' comprising of all place cell's firing rate-by-position vector as:

$$Pr_{(pos|spikes)} = (\prod_{i=1}^{n} f_i(pos)^{sp_i})e^{-\tau}\sum_{i=1}^{n} f_i(pos) \quad (4)$$

Where $f_i(pos)^{sp_i}$ is the value of the firing rate-by-position vector of the $i^{th}$ cell at position $pos$, $sp_i$ is the number of spikes fired by the $i^{th}$ cell in the time bin being decoded, $\tau$ is the duration of the time bin (100 ms) and *n* is the total number cells. Posterior probabilities were subsequently normalized to one:

$$Pr\ (pos|spikes)\ =\frac{Pr(pos|spikes)}{\sum_{i=1}^{p_n}Pr(pos_i|spikes)} \tag{5}$$

Where $p_n$ is the total number of position bins (5cm bins).

**Bayesian trial decoding.** Bayesian classification of trials was performed utilizing a template comprising of all cell's smoothed firing rate-by-pos-trial matrix from each arm as:

$$Pr_{(trial,pos|spikes)}\ =\ (\prod_{i=1}^{n}f_i(trial,pos)^{sp_i})e^{-\tau}\sum_{i=1}^{n}f_i(trial,pos) \tag{6}$$

Where $f_i(trial,pos)^{sp_i}$ is the value of the firing rate-by-trial vector of the $i^{th}$ cell at position *pos* and trial *trial*, $sp_i$ is the number of spikes fired by the $i^{th}$ cell in the time bin being decoded, $\tau$ is the duration of the time bin (100 ms) and *n* is the total number of place cells. Posterior probabilities were subsequently normalized to one using the *softmax* function:

$$Pr\ (pos|spikes)\ =\frac{e^{Pr(pos|spikes)}}{\sum_{i=1}^{p_n}e^{Pr(pos_i|spikes)}} \tag{7}$$

**Mixed-effect linear regression analysis.** To quantify the relative magnitude of the effects of different variables to explain the trial distribution pattern of pos-sleep replay, we applied mixed-effect linear regression analysis (MATLAB function fitlme). The model was fitted with the following formula:

$$post\ sleep\ trial\ replay\ pattern \sim maze\ replay\ trial\ pattern + theta\ cycle\ trial\ pattern + theta\ power\ trial\ patterh + pre\ sleep\ trial\ patter + shuffle\ trial\ pattern + (1|session_{ID}) \tag{8}$$

**Statistical analyses**. All statistical analyses were performed with custom-written scripts in MATLAB and Python. Power analysis was not used to determine sample sizes. Our study did not involve comparison of two or more groups; therefore, random allocation and experimenter blinding did not apply to our data.

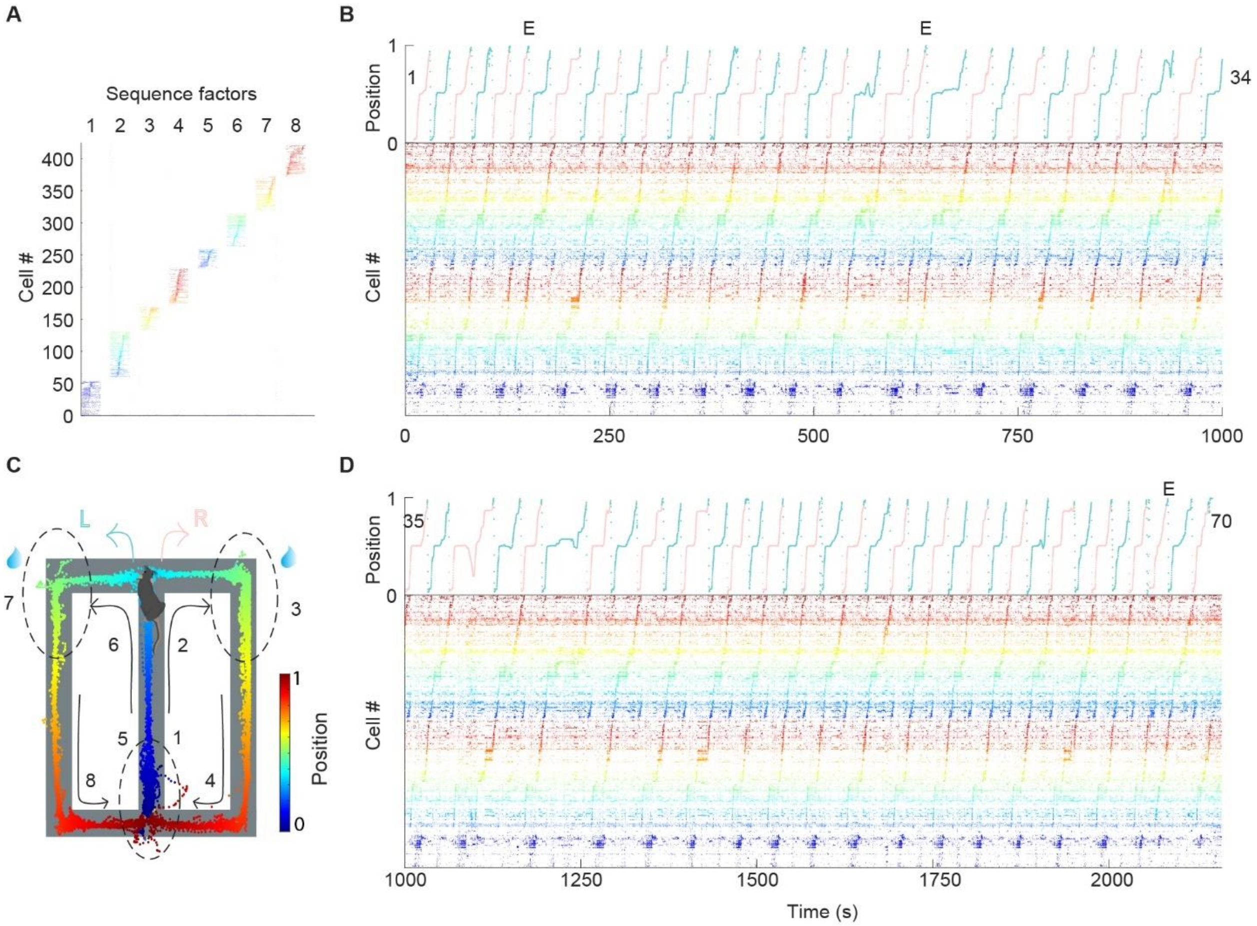


**Fig. S1: Neural spiking data of the entire example session sorted by seqNMF.**

**A.** 8 sequence factors identified by seqNMF. **B.** Trials 1-34 during the figure-8 maze task. Top, linearized position of the animal. Red, right traversal; green, left traversal. Bottom, raster plot of spiking data of 422 pyramidal cells simultaneously recorded from the right dorsal CA1 region, sorted by seqNMF, and color-coded according to the position (the same color scheme as in Fig. 1B). **C**. Figure-8 maze task, where mouse alternates between left and right arms to gain water reward (blue droplets). Animal's trajectory on the maze is colored by its linearized position. Numbers 1 to 8 indicate the mapping of the 8 seqNMF sequence factors onto different segments of the maze. **D.** Trials 35-70 of the same session as in B.

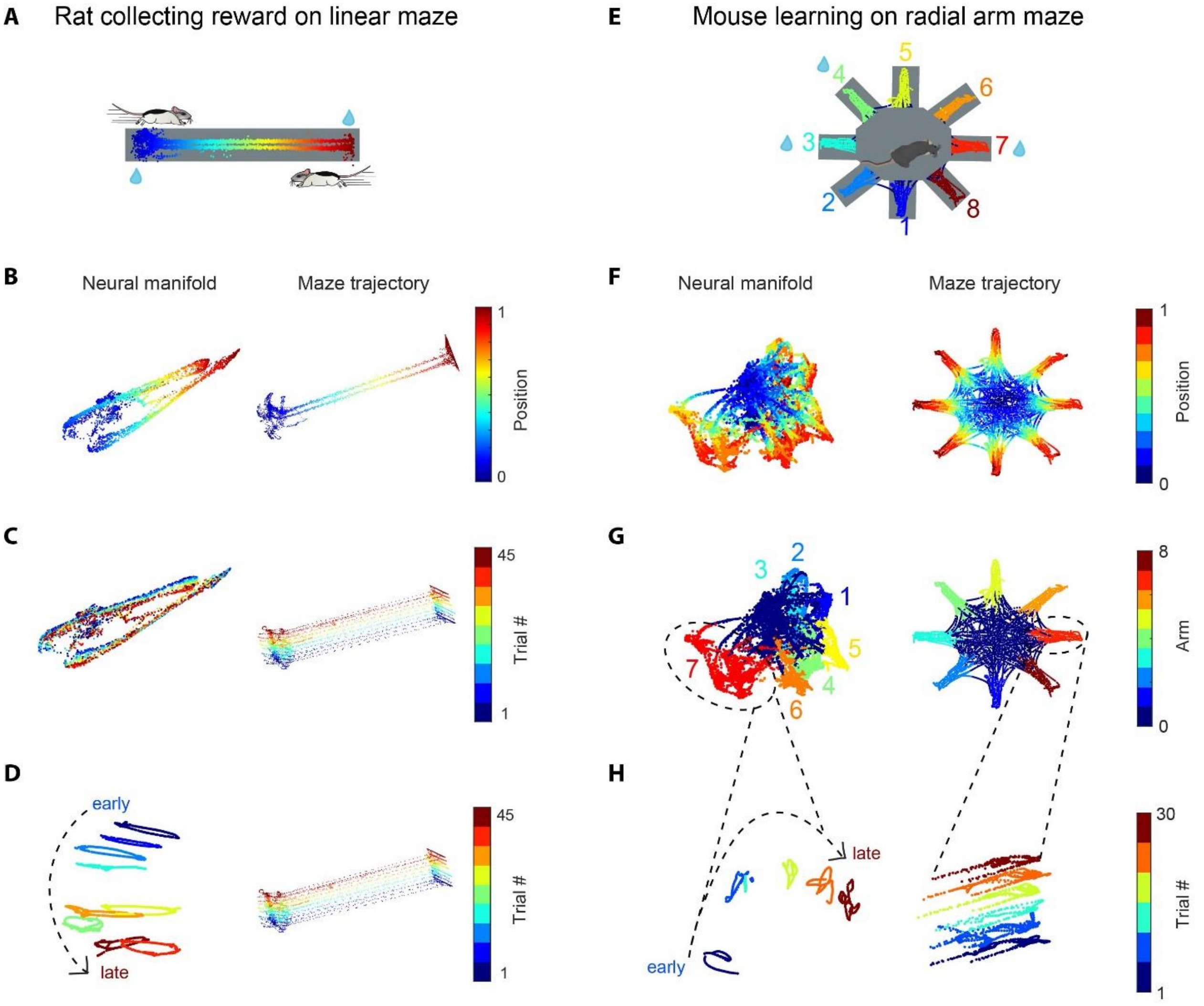


**Fig. S2: Trial-specific evolution of neuronal population activity can be observed in different tasks and rodent species**. Task topology and trial specific population representation from the manifold of figure-8 maze (Figure 1) were also observed in the manifolds of simpler tasks like linear maze, as well as more complex tasks like radial arm maze. **A.** Rat running back and forth between the two reward ports located at either end of the linear track to collect water reward (blue droplets). Animal's trajectory along the track is color-coded by animals' linearized position along the track. Only data when the animal's speed is larger than 1 cm/s is included. **B**. Left, UMAP embedding of population activity, colored by the rat's linearized position. Right, running trajectory of a rat on the linear maze. **C**. Same session as B but the manifold (left) and maze trajectory (right) are colored by trial block number. **D.** Same session as B, C, but using semi-supervised UMAP trained on blocks of 5 trials. The neural manifold (left) and maze trajectory (right) are colored by trial block number. **E.** In the radial-arm maze experiment, mouse collect water reward (blue droplets) from 3 out of the 8 potential reward sites at the end of each arm of the radial arm maze (arms 3,4 and 7 were rewarded in this example session). The animal's trajectory along the track is color-coded by the arm identity. **F.** Left UMAP embedding of population activity, colored by animal's linearized position on each of the 8 arms. Right, running trajectory of a mouse on the radial arm maze. **G**. Same session as F but the neural manifold and maze trajectory are colored by arm identity. 0 represents the center area. **H.** Since the manifold of figure-8 maze is complex, we focused only on one of the 8 arms to have a clear visualization of the state space evolution with trial number. We focused on arm 7, because the mouse visited this arm for 30 trials on this recording day. Neural manifold (left) was trained using semi-supervised UMAP on blocks of 5 trials. The manifold (left) and maze trajectory (right) are colored by trial block number.

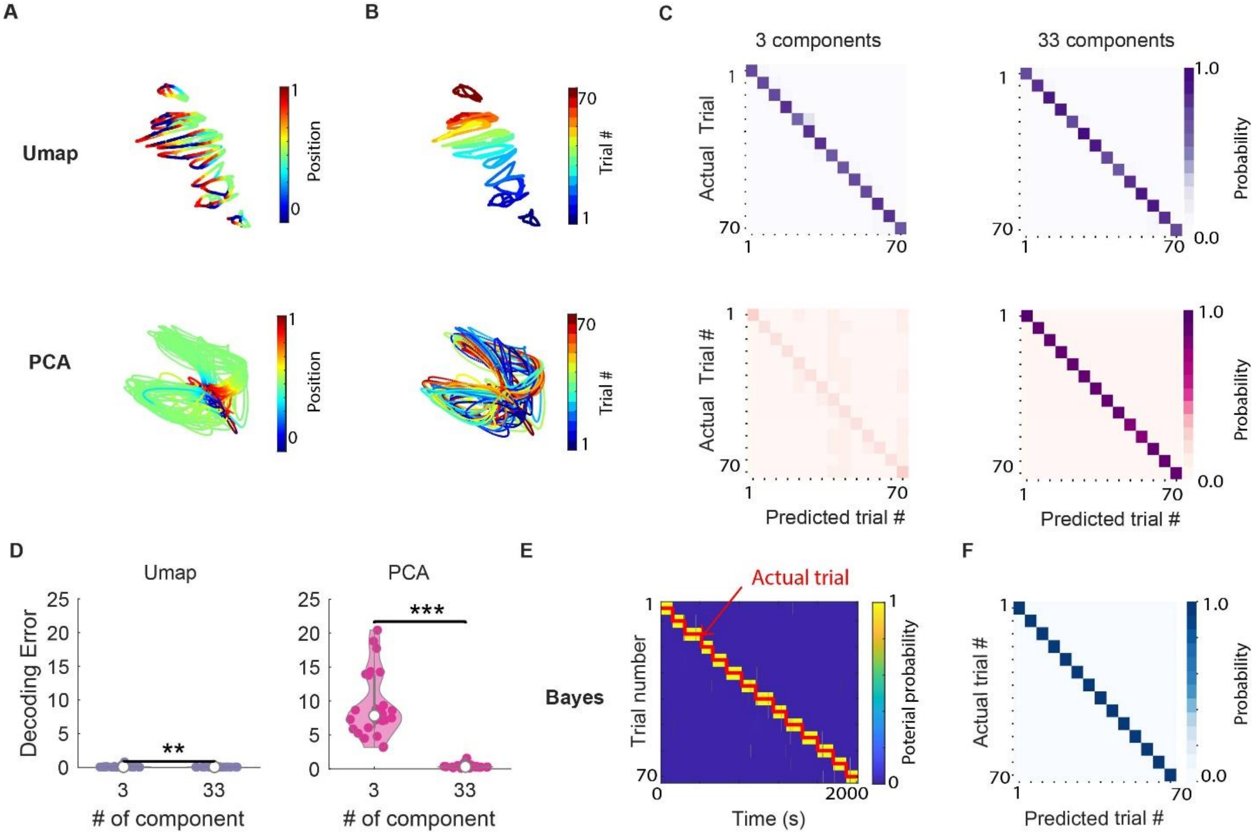


**Fig. S3: Trial identity can be decoded using three different methods.**

**A.** Low dimensional embedding of CA1 population activity, colored by animal's linearized position on the figure-8 maze. Top, UMAP manifold. Bottom, PCA manifold. **B.** Same as A but colored by trial block number. Top, UMAP manifold. Bottom, PCA manifold. **C.** Confusion matrix of trial decoding accuracy using 3 (left) and 33 (right) components of the low dimensional data. Top, UMAP decoding accuracy. Bottom, PCA decoding accuracy. **D.** Comparing decoding accuracy using UMAP and PCA using the first 3 and 33 components (n=26 sessions in 6 animals) of the low-dimensional space. PCA decoding achieved a similar level of decoding accuracy as UMAP when decoded based on the first 33 principal components. ** $P < 0.01$, *** $P < 10^{-9}$ for UMAP and PCA respectively; paired t-test; n = 26 sessions from 6 animals. **E**. Posterior probability of Bayesian trial decoding across time in one session. Red line, actual trial number. **F.** Confusion matrix of trial decoding accuracy from Bayesian decoding for the session in D.

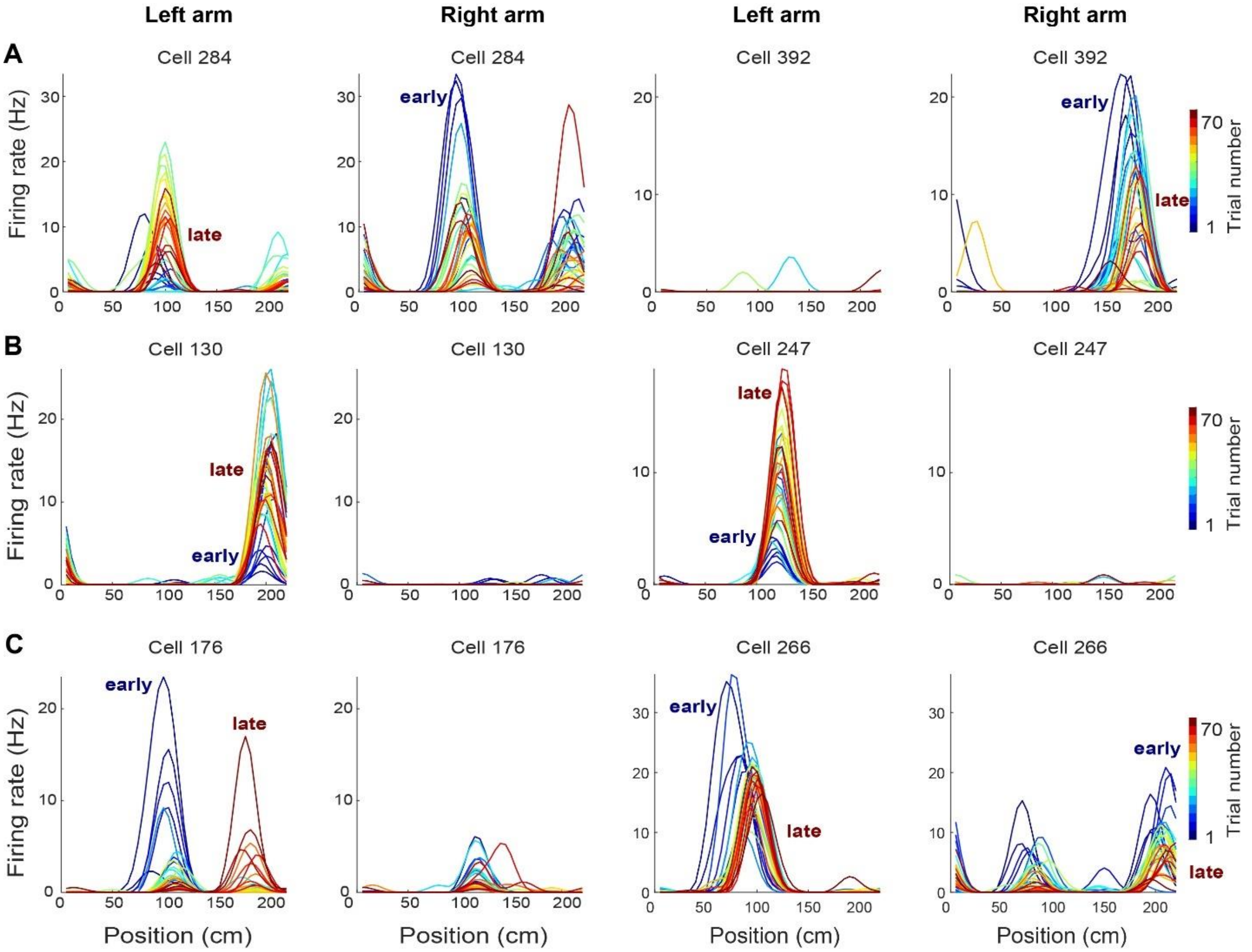


**Fig. S4: Tunning curve across trials of 6 example neurons.**

**A.** 2 example cells that fired with higher firing rate at early trials. Left, left arm trials. Right, right arm trials, **B.** 2 example cells that fired with higher firing rate at late trials. **C.** 2 example cells with place field remap across trials.

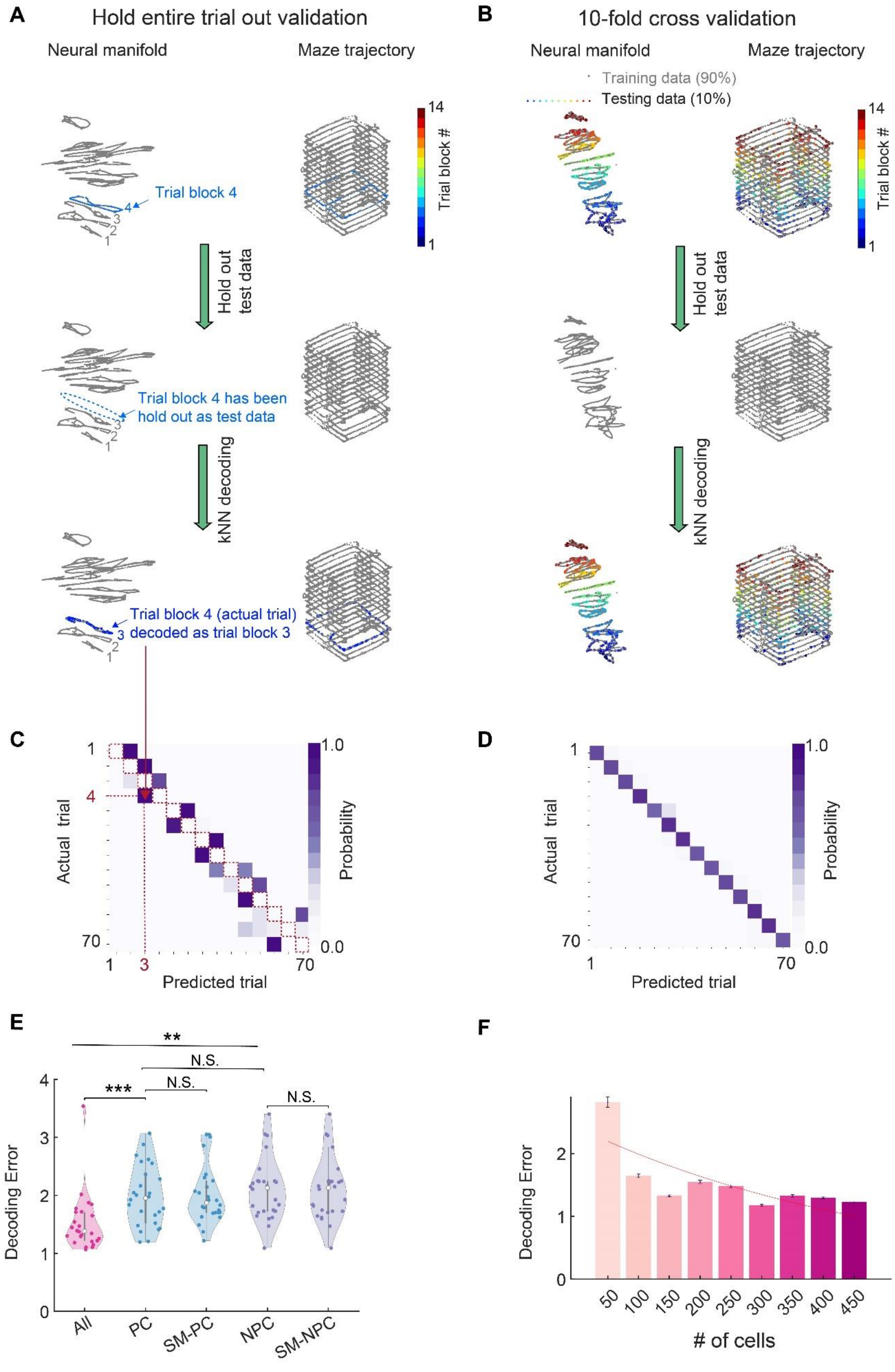

A
Hold entire trial out validation
Neural manifold
Maze trajectory
Trial block 4
Trial block #
Hold out test data
Trial block 4 has been hold out as test data
kNN decoding
Trial block 4 (actual trial) decoded as trial block 3
B
10-fold cross validation
Neural manifold
Maze trajectory
Training data (90%)
Testing data (10%)
Trial block #
Hold out test data
kNN decoding
C
Actual trial
Predicted trial
Probability
D
Actual trial
Predicted trial
Probability
E
Decoding Error
All
PC
SM-PC
NPC
SM-NPC
**
***
N.S.
F
Decoding Error
### of cells

**Fig. S5. Two validation methods.**

**A.** Illustration of the hold entire trial out validation method. Top, neural manifold from all the data in one example session. Data from one trial block (trial block 4; blue) is highlighted. Middle, the highlighted trial block in A was held out as test data. After removing the test data, a kNN decoder was trained only using training data (grey). Bottom, the test data was then projected onto the manifold and the kNN decoder was used for trial identity decoding. Training data is gray and test data is colored by the identity of the decoded trial. Test data was decoded to the trial block label of its nearest neighbor on the training data manifold. **B.** Illustration of the 10-fold cross-validation method. Top, neural manifold generated from all the data in one example session. 10% of the data are highlighted (colored by their true trial identity). The highlighted data is held out as test data. Middle, after removing the test data, a kNN decoder was trained only using training data (grey). Bottom, projecting the testing data onto the manifold and use KNN decoder for trial identity decoding. Training data is gray and test data is colored by the identity of the decoded trial (trial block 3). **C.** Confusion matrix obtained using hold entire trial out validation. Note that the diagonal (red dashed line) of the confusion matrix has 0 decoding probability, because the training and test data does not share data from the same trial block. Since the neural embedding was structured according to progression of trial events, the test data were decoded to their nearest neighbor. **D.** Confusion matrix obtained from 10-fold cross validation. **E**. Trial decoding error comparing all cells, place cells and their size-matched control (purple), as well as none-place cells and their size-matched control (blue). Here hold entire trial out validation was used. Decoding error was measured in unit of trial block (where successive 5 trials were binned to 1 trial block). *** $P < 0.001$, ** $P < 0.01$, N.S. not significant; unpaired t-test; N = 26 sessions from 6 animals. **F**. Decoding error when down-sampling the cell numbers in the example session from 450 to 50. Decoding error was measured in unit of trial block (where successive 5 trials were binned to 1 trial block). Error bar indicating the standard error of mean (SE) across 10 different random seeds that selected different random subsamples of cells. Red line, fitted exponential curve. Similar to Fig. 2J, where tenfold cross validation was used, and leave one-trial out validation was used here.

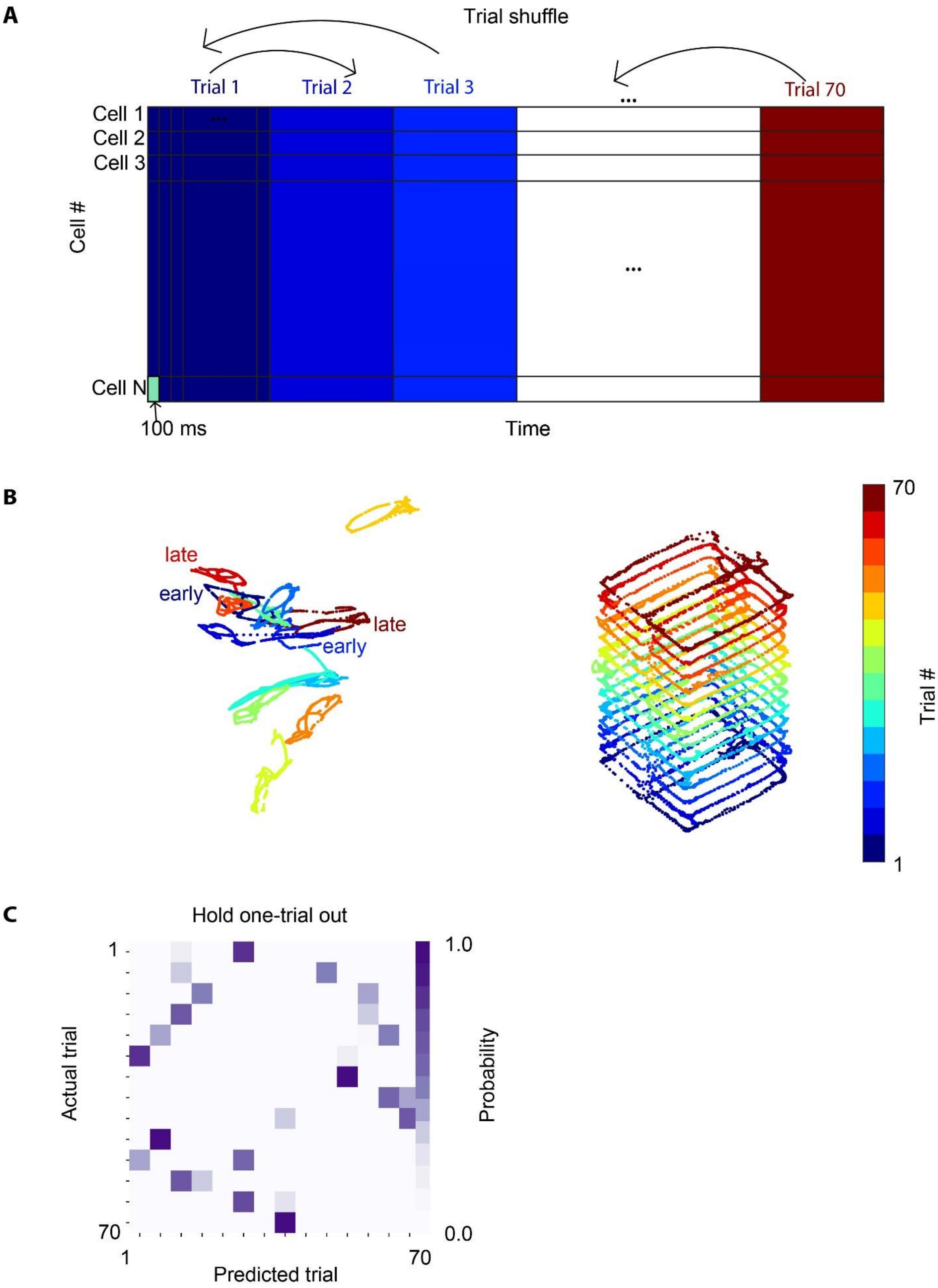
A
Trial shuffle
Trial 1
Trial 2
Trial 3
Trial 70
Cell 1
Cell 2
Cell 3
Cell N
Cell #
100 ms
Time
B
late
early
late
early
70
Trial #
1
C
Hold one-trial out
1
70
Actual trial
1
70
Predicted trial
1.0
0.0
Probability

**Figure S6. Trial shuffle procedure and its decoding results**

**A.** Illustration of the trial shuffle procedure. Neural spiking data was binned by 100ms bins and concatenated into a matrix (each row is a cell, and each column is a 100ms bin). Trial shuffle was done by shuffling the trial identity of the entire data matrix (same shuffle for all the cells). **B.** Left, neural manifold for the trial-shuffled data. The neural manifold (left) was colored by the shuffled trial identity and the maze trajectory (right) was colored by the true trial identity. Notice that the shuffle procedure disrupts the structure of the data such that early and late trials are embedded very close to each other. **C.** Confusion matrix of the trial shuffled data hold-entire-trial-out validation. Trial shuffled data lost the trial structure in real data -- trials close in event order were no longer embedded close in the state space. Thus, hold one-trial out validation method yields low decoding accuracy for trial shuffled data.

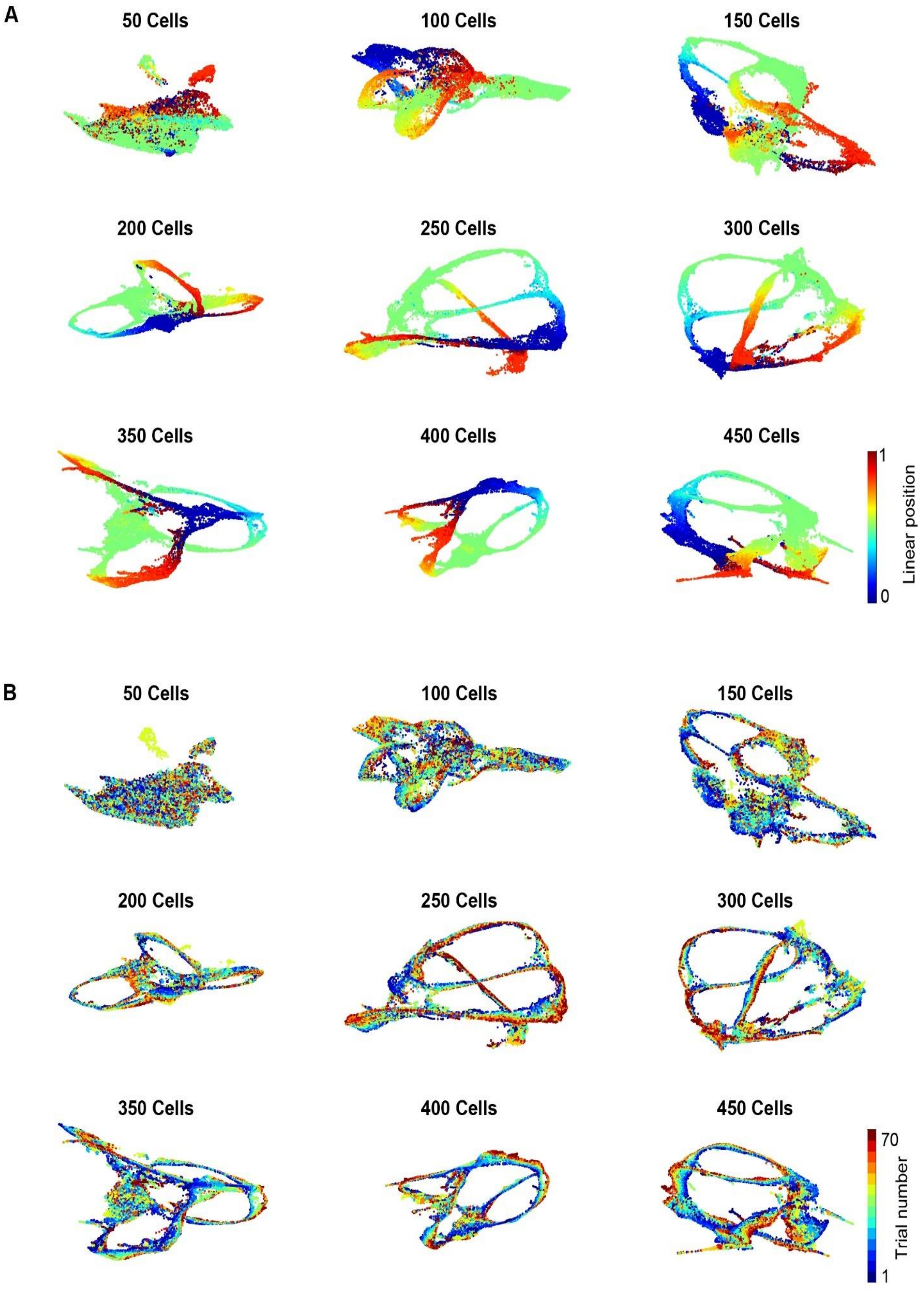


**Figure S7. Visualization of down-sample analysis using UMAP (unsupervised)**

**A.** Manifolds generated from down-sampling the number of cells used as input to UMAP. Manifolds were colored by animal's linearized position. **B.** Same as in **A** but manifolds were colored by trial number.

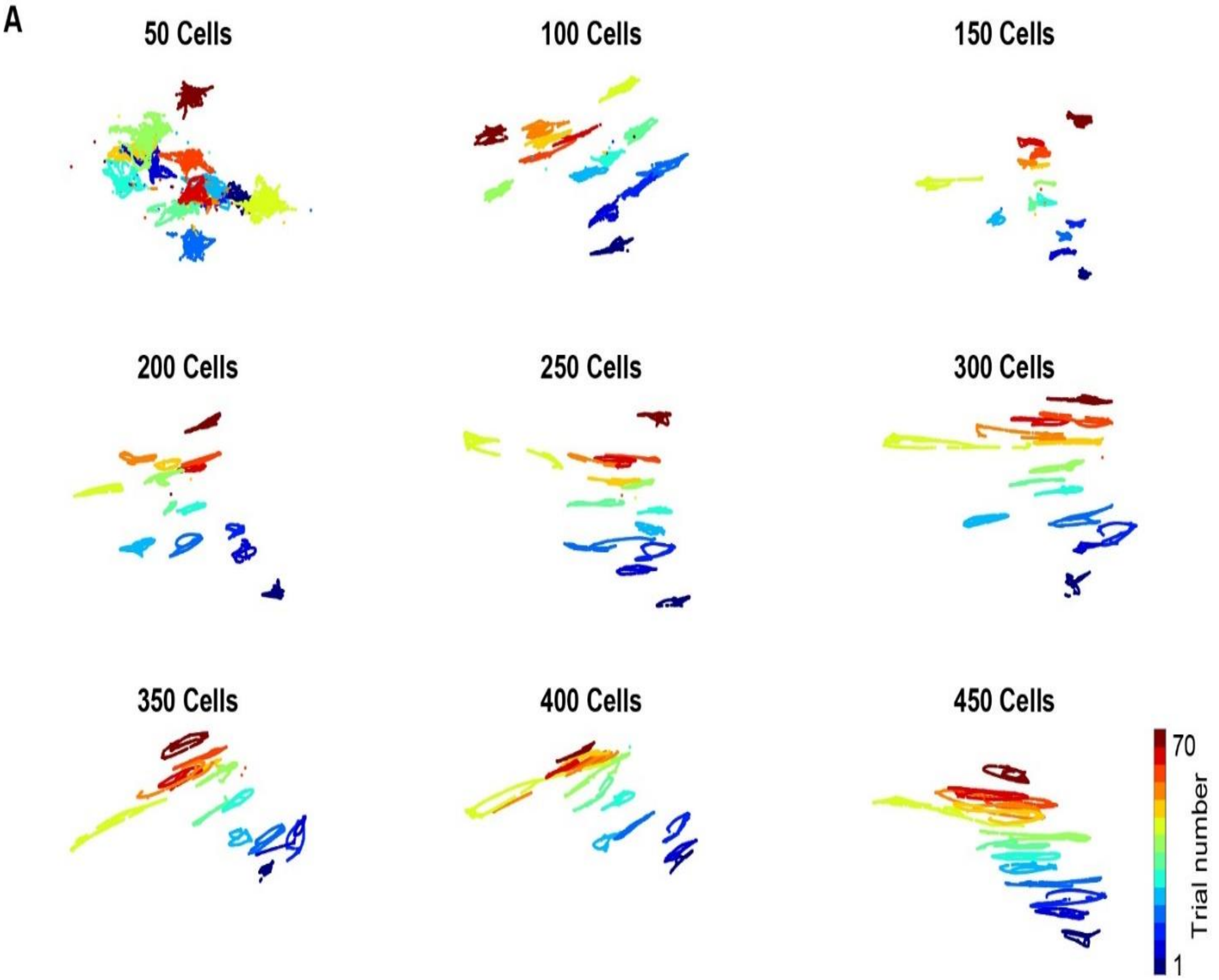


**Figure S8. Visualization of down-sample analysis UMAP (semi-supervised)**

**A.** UMAP manifolds generated from down-sampling the number of cells. Here, the manifold embeddings were trained by semi-supervised method. Manifolds were colored by trial block number.

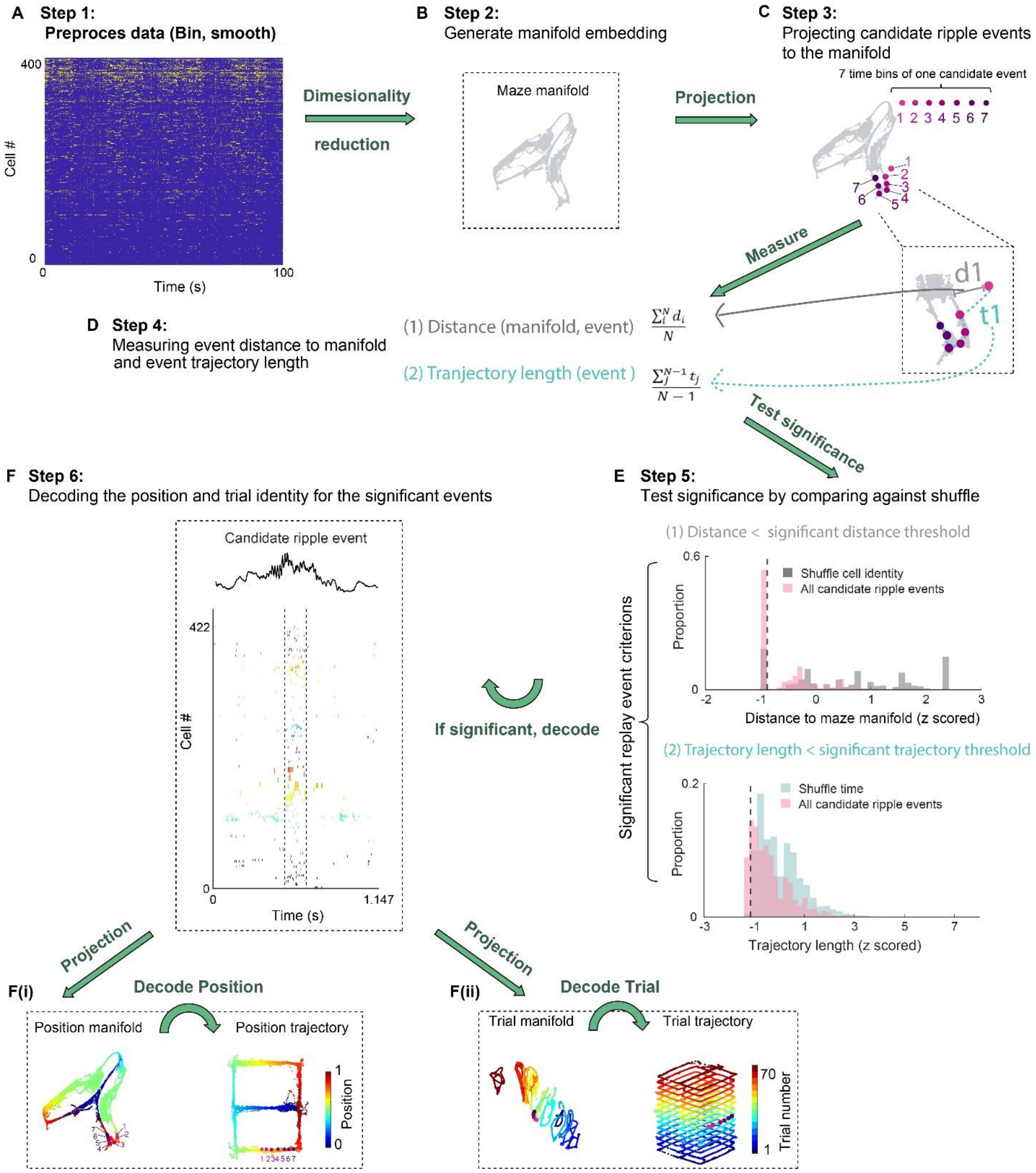


**Figure S9. Flow diagram demonstrating procedures for detecting and decoding significant SPW-R replays.** **A.** Step 1: Preprocessing data as follows. The animal performed a learning task in the figure-8 maze while spikes from 422 pyramidal cells shown in Fig. 1A were recorded (cells are ordered arbitrarily). Spike trains during maze learning were binned into 100ms bin and then smoothed, yielding a matrix of 422-dimensional population activity vectors. Spike trains of candidate SPW-R replays were binned into 20ms bins. **B.** Step 2: generating manifold embedding. The preprocessed data matrix is passed through dimensionality reduction step by either UMAP or PCA, which generates a low dimensional embedding, allowing the topological structure to be visualized. Notice that this step is un-supervised. For further details related to step 1 and 2, see Suppl. Fig. 10. **C.** Step 3: Projecting candidate ripple events to the manifold. The 7 pink-purple dots on the manifold represent low-dimensional embeddings of an example candidate SPW-R event. Each dot is the low-dimensional embedding of a 20ms bin. **D.** Step 4: measuring distance to manifold and trajectory length (as in Fig. 3D). The event distance to manifold is defined as the mean Euclidian distance to manifold across all time bins within the event. For example, grey bar in the zoomed-in inset represents the distance between the first time bin and the maze manifold (d1). The event trajectory length is defined as the mean trajectory length between all successive time bins. For example, cyan dashed line denotes the trajectory length between the first and second time bin of the event (t1). **E.** Step 5: Testing whether the candidate events are significant by comparing them against shuffled distributions. A candidate event is classified as a significant replay event only if it satisfies two criteria: (1) The mean distance to manifold (as defined in D) of a candidate event is significantly smaller than shuffled data (shuffle cell identity). (2) The second criterion is that the mean trajectory length is significantly smaller than shuffled data (shuffle time). See Suppl Fig. 11 for further explanation of how the shuffle distributions were generated**.** **F.** Step 6: Only significant replays are included for further analysis from this step onward. **F(i).** To decode the position relayed by the candidate event, the population activity during SPW-R was projected to the position manifold where each point on the manifold is associated with a position bin label (positions were binned into 5-cm bins). The low dimensional embedding along with the position label are used to train a KNN decoder. Replay content of each time bin is decoded by taking the mean of the position label of its K nearest neighbors on the position manifold. **F(ii).** Similarly, the SPW-R activity vectors are projected to the trial manifold for trial identity decoding. If using UMAP, the trial manifold is learned through semi-supervised dimensionality reduction (trials are binned into 5-trial bins). The similar procedure can be applied for decoding position and trial content of SPW-R events using PCA.

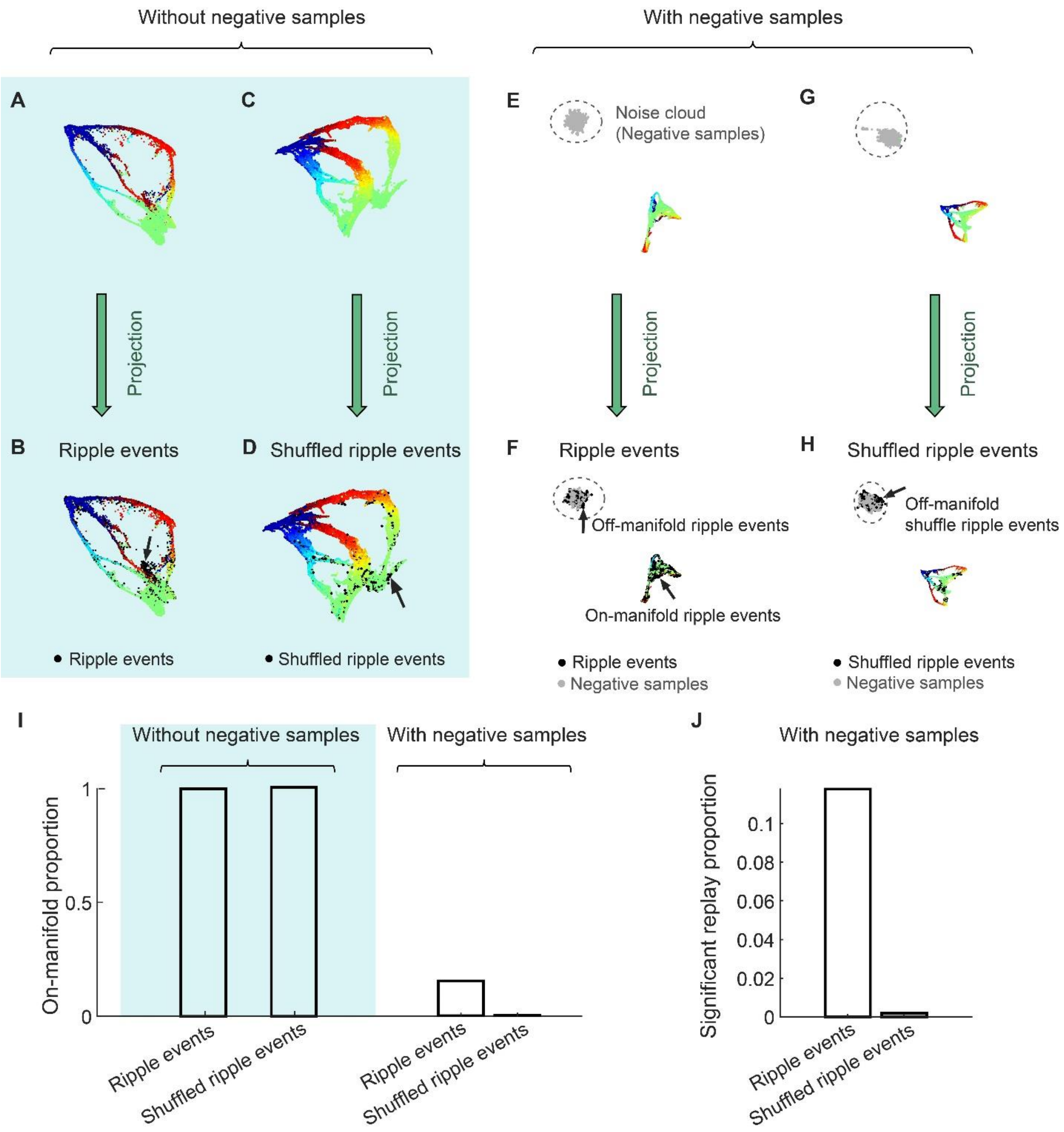


**Figure S10. Metric learning and negative sampling.**

This figure is paired with Step 2 and 3 of the decoding procedure described in Suppl Fig. S9. Although UMAP is typically used as a self-supervised dimensionality reduction tool. It can also be used to do metric learning by embedding out-of-sample points based on an existing embedding. Briefly, metric learning refers to a process where a labeled set of points is used to learn a metric on data, and then the learned metric is used as a measure of distance between new unlabeled points. Since SPW-R data is unlabeled and maze data is labeled (labeled by behavior-related variables like position and trial identity, etc.), metric learning provides a natural framework for decoding the content of candidate replay events. **A.** To use metric learning for decoding, UMAP embedding from maze data is generated. The learned metric is used as a measure of distance between new unlabeled points. **B.** Second, population vectors during ripple events are projected to the maze manifold. Note that almost all SPWR-R events are embedded onto the maze manifold. **C.** and **D**. are similar to A and B, but here shuffled SPW-R events are projected to the manifold. Even shuffled events are embedded on the maze manifold. This happens because when optimizing the low-dimensional embedding, the UMAP algorithm uses negative sampling, which attracts the positive sample pairs and repulse the negative sample pairs (*3*, *58*). This explains why even shuffled data that are supposed to be far from manifold are embedded onto the manifold: in the absence of negative samples, there is no repulsive force and so every data point is mapped to the maze manifold. **E**. To avoid collapsed embedding, negative samples (grey dots) are generated and embedded along with the maze manifold so that noise is attracted to the negative sample cloud. **F**. Only SPW-R events (black), which are similar to the maze manifold are mapped onto the maze manifold. SPW-R events that are different from maze manifold are attracted to the noise cloud (circled). **G**. and **H** are similar to **E** and **F**, but instead of projecting real SPW-Rs events, shuffled events are projected to the learned embedding metric. **I**. Without negative samples, all SPW-Rs events and even shuffled data are on-manifold. In contrast, with negative samples, only 15% of real SPW-Rs events and 0.2% of shuffled events are on-manifold. **J**. For an event deemed to be significant replay event, it needs to be not only on-manifold (small distance to manifold comparing to shuffle), but also has a short trajectory length (comparing to shuffle). See step 4 in Suppl. Fig. 9, as well as Fig. S11. Approximately 11% of the SPW-R events are significant replay events, whereas 0.1% of the shuffle events satisfy both significance criteria.

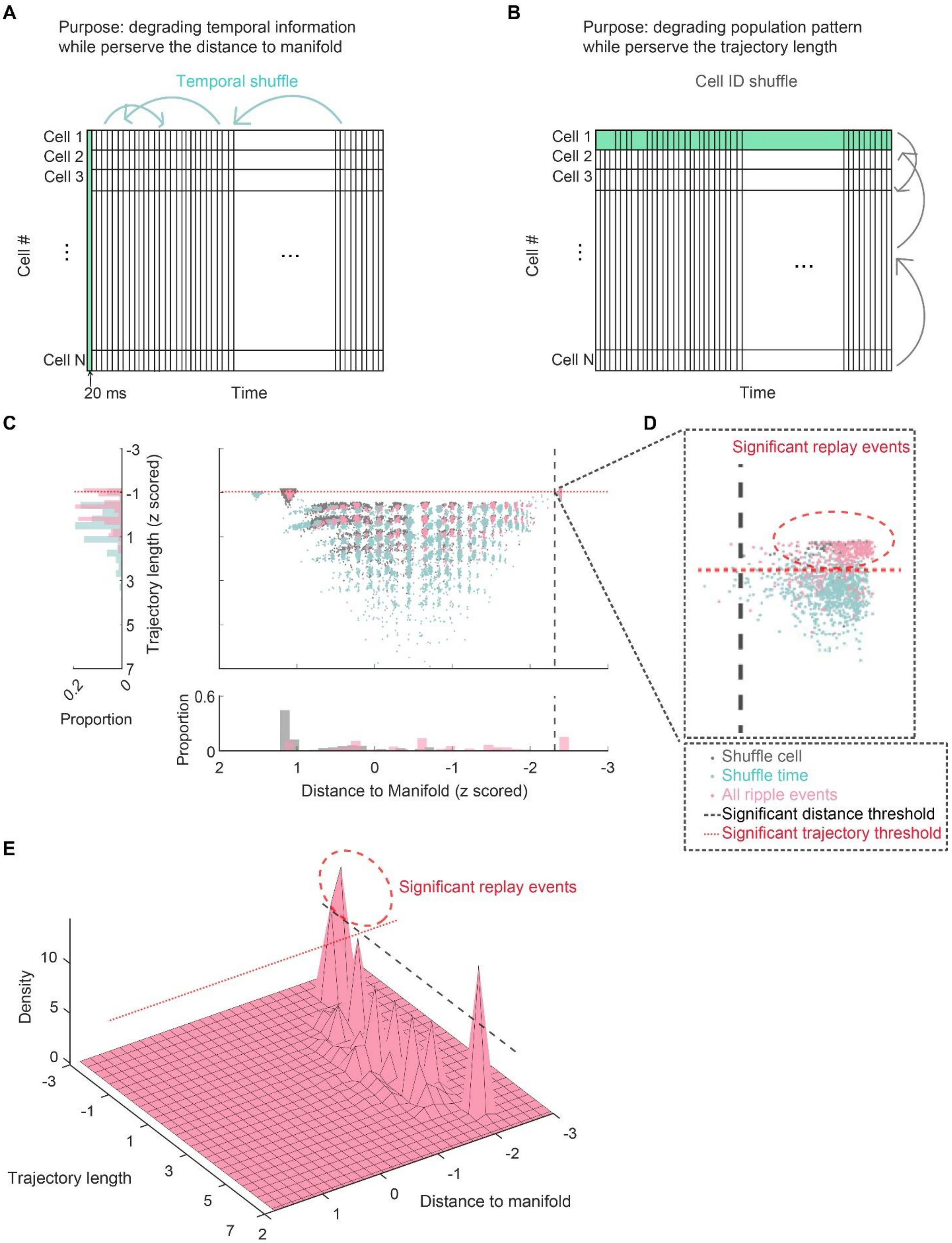


**Figure S11. Quantifying significant SPW-R replay events against shuffle events.**

This figure is related to Step 5 of the decoding procedure described in Suppl Fig. S9. Candidate SPW-R events are classified as significant replay events only if the distance to manifold and trajectory length are significantly small compared to null distributions. **A.** Shuffling procedure for degrading the temporal information of candidate SPW-R events without changing other properties of population activity pattern (preserving relationship across rows). This is done by shuffle cell ID. Spiking activity of each cell is a row of the matrix, and each column is a 20ms bin. This procedure is expected to generate shuffled events that have a long trajectory length compared to true replay events without affecting their distance to the manifold. **B.** Temporal shuffle. This procedure degrades the population activity pattern while preserving the temporal information of SPW-R events. This is done by shuffling the rows of the spiking matrix (by preserving relationship across columns). This procedure is expected to generate shuffled events that have large distance from manifold compared to true replay events, without affecting their trajectory length. **C.** Middle, scatter plot of events spreading in a 2-D metric defined by the z-scored distance to manifold and trajectory length. Bottom, histogram for distance to manifold distribution comparing the shuffled data (grey, shuffle cell as in B.) and all SPW-R events (red) in an example session. Left, histogram for trajectory comparing the shuffled data (cyan, shuffle time as in A) and all SPW-R events (red). **D.** Zoomed-in display significant replay events. Black dash line, significant distance threshold. Red dash line, significant trajectory threshold. **E.** For better visualization of the distribution of SPW-R events, density plot is shown, where the density value is the z-scored count of events in each of 30x30 grid of the distance and trajectory metric. Black dash line, significant distance threshold. Red dash line, significant trajectory threshold.

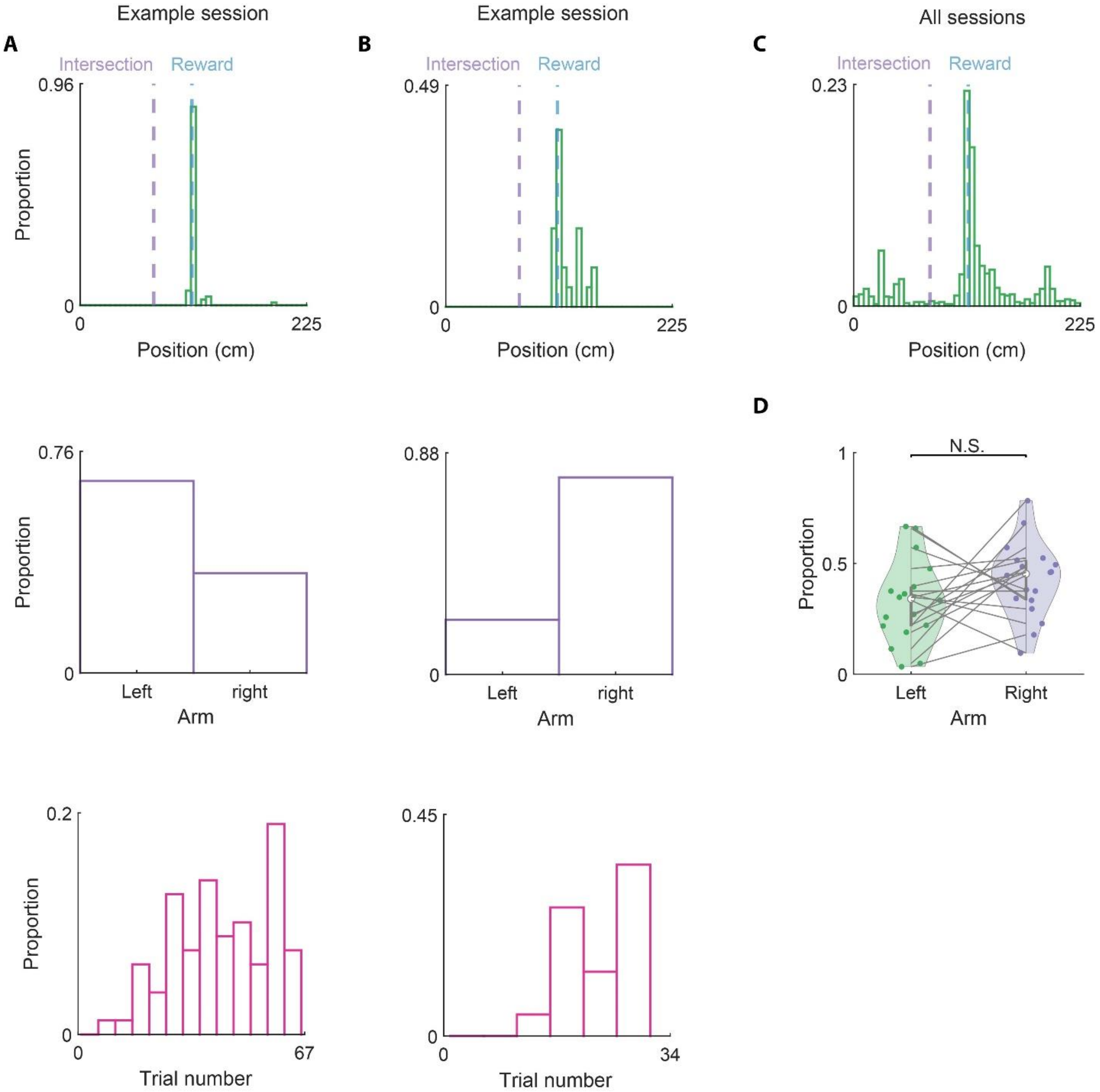


**Figure S12. Distribution of SPW-Rs across different locations, trials, and arms of the maze.**

**A-B.** Distribution of SPW-Rs across different maze locations (top), trials (middle) and arms (bottom) of the figure-8maze from two example sessions. Intersection, T junction of the figure-8 maze. Reward, location of the water reward in the left or right-side arm. **C.** Distribution of SPW-Rs across different locations pooled from all figure-8 maze sessions; n = 21 sessions from 6 animals. **D.** Distribution of SPW-Rs across different arms of the maze, pooled from all figure-8 maze sessions. N.S., not significant (P = 0.11); unpaired t-test; n = 21 sessions from 6 animals.

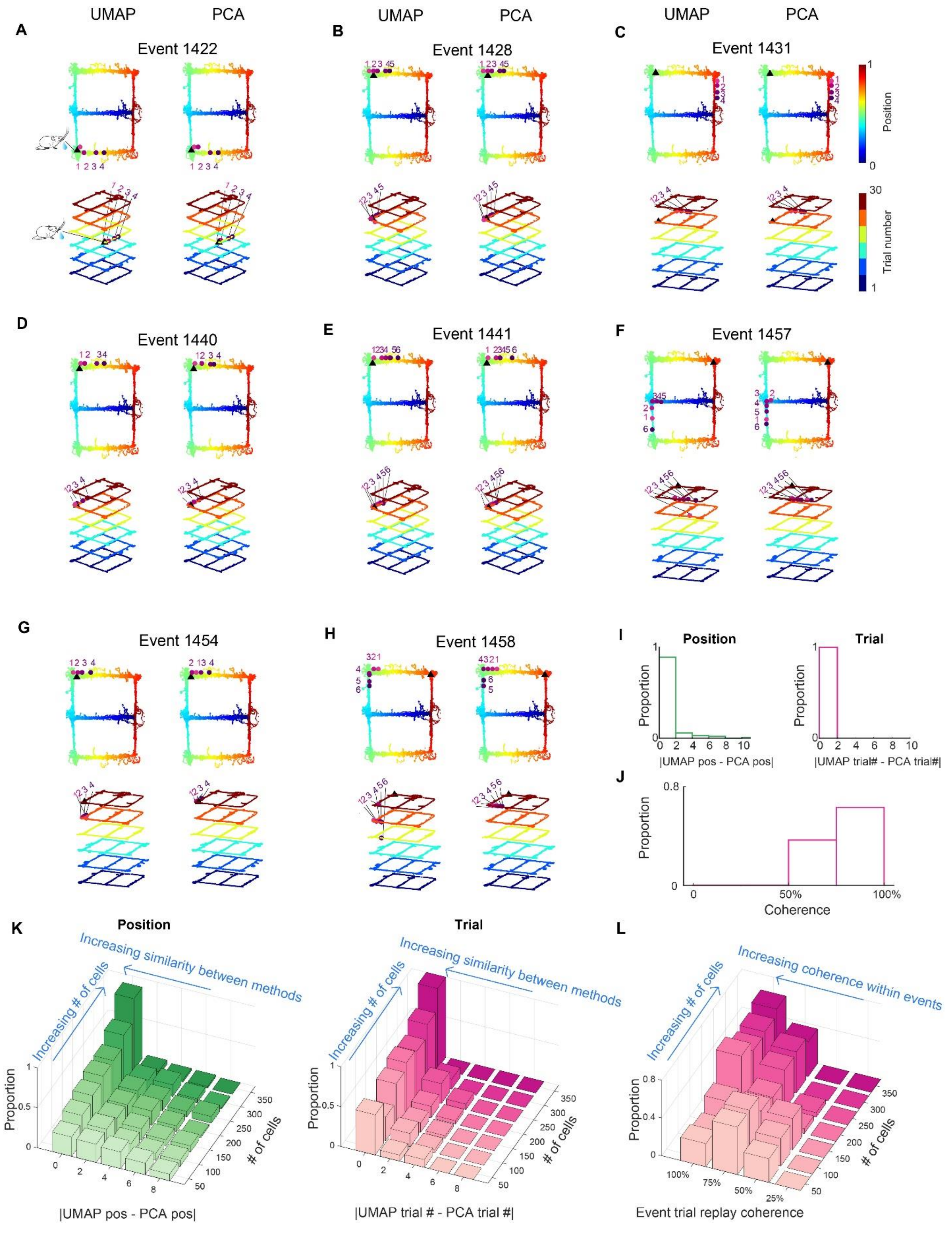


**Figure S13. Matching between UMAP and PCA and coherence of trial replay.**

**A-H.** Example replay events from an example session. For each event, the left columns are UMAP decoding results, and the right columns are PCA decoding results. The top rows are position decoding results and bottom rows are trial decoding results. **I.** Matching between UMAP and PCA decoding results, measured by the difference in decoding results from the two methods. Left, matching between UMAP and PCA results for position decoding (measured by absolute value of the difference between UMAP decoded position bin and PCA decoded position bin). Right, matching between UMAP and PCA results for trial decoding (measured by the absolute value of the difference between UMAP decoded trial block and PCA decoded trial block). 0 means PCA and UMAP decoded to the same position bin or trial block. For example, event 1422 in panel A has completely coherent decoding results between PCA and UMAP decoding (0 difference). **J.** Event trial replay coherence distribution. The coherence of each event was measured as the proportion of time bins that share the same trial label as the mode for that event. 100% means all time bins of a given replay event are decoded to the same trial label. For instance, event 1422 in panel A is an example even with 100% coherence (all 4 time bins decoded to trial block 4). **K.** One potential issue reported in the field is that different decoding methods produce different decoding results (59). We hypothesized that the low decoding coherence between different methods reported in earlier studies may result from the limited number of neurons available for analysis. To test this hypothesis, we down-sampled neurons used for the decoding analysis. Left, matching between UMAP and PCA for position decoding. Right, matching between UMAP and PCA for trial identity decoding. Note that UMAP and PCA produce similar results (0 difference, first column) when the number of neurons used for decoding is high but begin to diverge with a decreasing number of neurons. **L.** As in K, we performed down-sample analysis to decrease the number of neurons used for decoding. More neurons used for decoding results in higher trial replay coherence (100% coherence means all the time bins of the event decoded to the same trial).

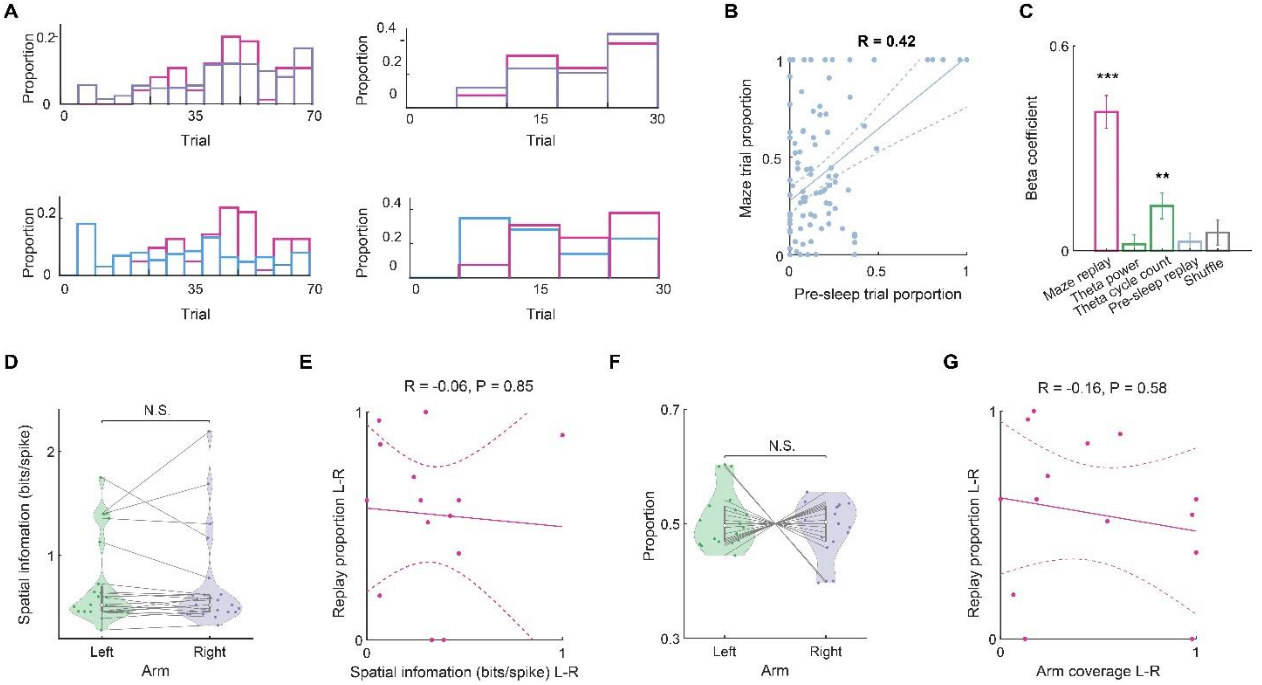


**Figure S14. Control analyses for trial and arm distribution**

A. Similar to Fig. 4G, distribution of the replay events across trials decoded from the population spike content of SPW-Rs in two example sessions. Top, maze and pos-sleep replay trial distribution pattern. Bottom, maze and pre-sleep replay trial distribution pattern. **B.** Correlation between trial distributions during maze VS. pre-sleep replay. Pearson correlation coefficient, $R = -0.16$, $P < 10^{-5}$. **C.** Similar to Fig. 4I but the results here were obtained from PCA decoding. *** $P < 10^{-12}$ for awake replay, ** $P < 0.01$ for theta cycle number; A variable could not significantly explain the target if the 95% confidence interval of its beta coefficient does not overlap with 0. **D.** Mean spatial information for left versus right arm segments. Each dot in the violin plot represents mean spatial information across all cells from one session. N.S. not significant, P = 0.76; unpaired t-test; n = 13 sessions from 5 animals. **E.** Correlation between spatial information difference (mean spatial information of left arm trials minus mean spatial information of right arm trials) and replay proportion difference between left and right arm (proportion of left arm replay minus proportion right arm replay). Pearson correlation coefficient, R = 0.07, P = 0.85; n = 13 sessions from 5 animals. This shows the difference in replay distribution across arms could not be explained by the difference in decoding quality across arms. **G.** Proportion of left versus right arm visit. Each dot in the violin plot represents one session. N.S. not significant; P = 0.33; unpaired t-test; n = 13 sessions from 5 animals. **G**. Correlation between the difference in arm visit (proportion of left arm visits minus proportion of right arm visits) and difference in replay proportion between left and right arm (proportion of left arm replay minus proportion right arm replay). Pearson correlation coefficient, R = -0.16, P = 0.58; n = 13 sessions from 5 animals.
